# Labile assembly of a tardigrade protein induces biostasis

**DOI:** 10.1101/2023.06.30.547219

**Authors:** S. Sanchez-Martinez, K. Nguyen, S. Biswas, V. Nicholson, A.V. Romanyuk, J. Ramirez, S. KC, A. Akter, C. Childs, E.T. Usher, G.M. Ginell, F. Yu, E. Gollub, M. Malferrari, F. Francia, G. Venturoli, E.W. Martin, F. Caporaletti, G. Giubertoni, S. Woutersen, S. Sukenik, D.N. Woolfson, A.S. Holehouse, T.C. Boothby

## Abstract

Tardigrades are microscopic animals that survive desiccation by inducing biostasis. To survive drying tardigrades rely on intrinsically disordered CAHS proteins that form gels. However, the sequence features and mechanisms underlying gel formation and the necessity of gelation for protection have not been demonstrated. Here we report a mechanism of gelation for CAHS D similar to that of intermediate filaments. We show that gelation restricts molecular motion, immobilizing and protecting labile material from the harmful effects of drying. *In vivo*, we observe that CAHS D forms fiber-like condensates during osmotic stress. Condensation of CAHS D improves survival of osmotically shocked cells through at least two mechanisms: reduction of cell volume change and reduction of metabolic activity. Importantly, condensation of CAHS D is reversible and metabolic rates return to control levels after CAHS condensates are resolved. This work provides insights into how tardigrades induce biostasis through the self-assembly of CAHS gels.

## Introduction

Water is required for all metabolism and is considered essential for life. However, a number of desiccation-tolerant organisms spread across the kingdoms of life challenge this thinking by surviving the loss of essentially all the hydrating water inside their cells. To perform this feat, desiccation-tolerant organisms enter into a state of biostasis known as anhydrobiosis (from Greek for “life without water”) [1]. In this anhydrobiotic state, organisms become ametabolic and remain so until rehydrated [1]. Once rehydrated, even after years or in some cases decades, these anhydrobiotic organisms resume their normal metabolism and development [1].

Many anhydrobiotic organisms protect their cells from drying-induced damage by accumulating non-reducing sugars such as sucrose [2] and trehalose [3–7]. The enrichment of disaccharides was long thought to be a universal feature of desiccation tolerance. However, several robustly anhydrobiotic organisms, such as tardigrades and rotifers, do not accumulate high levels, or in some cases any of these sugars during drying [8–10]. Instead, these animals use a diverse array of intrinsically disordered proteins (IDPs) to provide adaptive protection against desiccation [9,11–14].

One example of desiccation-related IDPs are Cytoplasmic Abundant Heat Soluble (CAHS) proteins [9,15–17], which are employed by tardigrades to survive drying [9,14,16,17]. Genes encoding CAHS proteins are induced in response to desiccation in tardigrade species that require preconditioning to survive drying, or are constitutively expressed at high levels in tardigrade species that require little priming to tolerate water loss [9]. Here, we focus on CAHS D, a representative CAHS protein derived from the tardigrade *Hypsibius exemplaris*, which is required for anhydrobiosis in this animal [9]. This protein increases desiccation tolerance when heterologously expressed in yeast and bacteria, and is sufficient to protect sensitive biological material from desiccation-induced damage *in vitro* [9,14,16]. CAHS proteins have been shown to be disordered through disorder prediction algorithms, solution NMR, hydrogen-deuterium exchange experiments, and circular dichroism spectroscopy [9,17].

Interestingly, several groups have observed that some desiccation-related IDPs, including CAHS D, can undergo a phase transition from solution to solid state, forming hydrogels. The gelation of CAHS proteins has been proposed to play a role in the protective capacity of CAHS proteins during desiccation [18–21]. However, sequence features that underlie this process have not been tested empirically, and the proposed molecular mechanisms that could underlie how CAHS proteins form gels are in conflict. Furthermore, the necessity of gelation in mediating stress tolerance has not been tested. Understanding if gelation is linked to stress tolerance and how gelation occurs in these proteins will provide insights not only into how CAHS proteins, and potentially other IDPs, undergo phase transitions, but will also provide avenues to further study of how CAHS proteins protect tardigrades during desiccation.

Here we demonstrate that CAHS D forms concentration- and temperature-dependent gels *in vitro* and that CAHS D forms gel-like condensates *in vivo* during osmotic stress. CAHS D gelation restricts molecular motions of CAHS D itself as well as those of desiccation-sensitive biological materials embedded within the gel. Using electron microscopy (EM) and small angle X-ray scattering (SAXS), we characterize the structure of CAHS D hydrogels and find that they are composed of a reticular network of ∼10 nm fibers. Using a combination of computational and biophysical approaches to characterize CAHS D, we show that this protein exists in a dumbbell-like ensemble consisting of two collapsed terminal regions rich in transient β-structure that are separated by a ‘Linker Region’ that forms an amphipathic α-helix. We find that while each region is necessary, none alone are sufficient to drive the solution-to-gel phase transition of CAHS D, suggesting that the protein’s dumbbell-like ensemble is essential for gelation. Using computational, biophysical, and molecular analysis, we arrive at a model of gelation where the amphipathic Linker Region drives bundling of CAHS D via helix-helix interactions, with higher-order networks assembling via β-β interactions between the terminal regions. We find that upon osmotic stress CAHS D forms condensates *in vivo*. Furthermore, CAHS D variants that perturb or modulate gelation *in vitro* also perturb or modulate condensates *in vivo*. During osmotic stress in cells, CAHS D condensation is required for resisting volume change, reducing metabolism, and improving survival. Interestingly, reduced metabolism during stress correlates strongly with increased survival rates. The condensation of CAHS D as well as its associated reduction of metabolism are reversible with the return to normal osmotic conditions. Thus, our data demonstrate that CAHS D forms a novel gel-like cytoskeletal assembly that promotes cellular survival during osmotic stress through the induction of reversible biostasis.

## Results

### CAHS D Forms a Concentration Dependent Reversible Gel

CAHS D (Uniprot: P0CU50) is a highly charged 227-residue protein that is disordered [9,17] and is required for the tardigrade *H. exemplaris* to survive desiccation robustly [9]. Through the course of studying this protein, we and others have observed that CAHS proteins undergo a solution-to-gel phase transition (Fig. 1A) [18,19].

**Figure 1.**
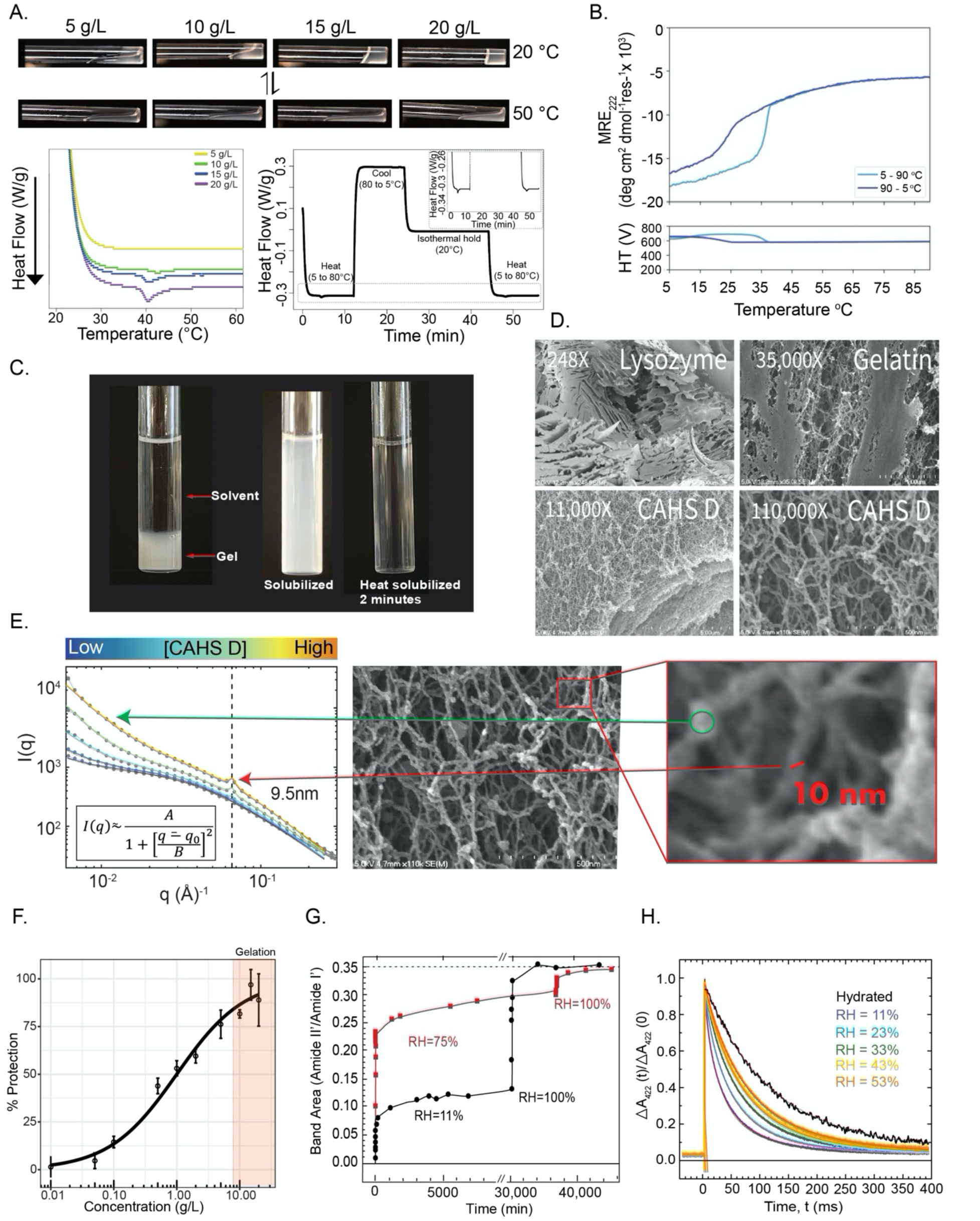
CAHS D forms a gel that immobilizes and stabilizes biological material *in vitro*. **A)** Concentration and temperature dependence of CAHS D gelation. CAHS D at 20°C (top panel) shows strong gelation at 20 g/L (0.8 mM), while at 50 °C gelation is disrupted. Once cooled, the gel reforms (lower panel right). Differential scanning calorimetry shows a concentration-dependent gel melt at ∼40 °C (lower left), which is reversible (lower right). **B)** Thermal denaturation and cooling of CAHS D (0.7 mM, 17.7 g/L) shows a sharp and highly reversible change in mean residue ellipticity (MRE), indicating the dissociation (heating, light blue line) and reassociation (cooling, dark blue line) of the protein. **C)** Dilution of 10 g/L (0.4 mM) CAHS D gel in 20 mM Tris buffer results in resolvation within ∼2 min at 55°C. **D)** SEM images of lysozyme, gelatin and CAHS D. CAHS D reticular gels are similar to the gel-forming protein, gelatin, but distinct from that of the non-gelling protein lysozyme. All SEM imaging was performed with proteins at 50 g/L. **E)** Probing CAHS D gels at various concentrations with SAXS revealed an emergent structure of approximately 9.5 nm. The region of the SEM image magnified in red callout box shows a fiber of approximately 10 nm in width, corresponding well with the emergent peak found at increasing concentration in the SAXS plot (indicated by the red arrow and black dotted line). The green arrow highlights the change in the internal void space within a single fiber. See also Fig. S1A & S1C. **F)** Plot of lactate dehydrogenase (LDH) enzyme stabilization by CAHS D showing the percent protection of LDH by CAHS D as a function of concentration. *n* = 3, Error bars = standard deviation. **G)** Kinetics of amide FTIR HDX in dried CAHS D matrices. Dried CAHS D matrices, equilibrated at RH = 6% were exposed to a D_2_O atmosphere at RH = 11% (black) or RH = 75% (red). Subsequently, both were transitioned to pure D_2_O atmosphere (RH = 100%), which demonstrated that both samples showed similar HDX when fully saturated. The dashed line represents the value of the amide II’ band area normalized to the area of the amide I’ band of CAHS D in D_2_O solution. **H)** Hydration-dependent immobilization of a client protein within the CAHS D glassy matrix. Charge recombination kinetics after a light pulse (fired at t = 0) are shown for photosynthetic reaction centers in solution (black) and embedded into dried CAHS D gels held at varying humidity. Colored traces represent different relative humidity (purple, RH = 11%; blue 23%; green 33%; yellow, 43%; orange 53%). Continuous red curves represent best fit to equation 1 (see Methods).

To characterize the material properties of CAHS D further, we assessed the concentration dependence of gelation. Using direct observation and differential scanning calorimetry (DSC), we found that CAHS D gelation is concentration dependent, with solutions below ∼10 g/L (0.4 mM) remaining diffuse, increasing in viscosity between 10 g/L - 15 g/L, and above ∼15 g/L (0.6 mM), forming robust gels (Fig. 1A). In addition to concentration dependence, gelation of CAHS D was observed to be reversible via heating and cooling (Fig. 1A-C) as well as by dilution (Fig. 1C). Reversibility suggests that gelation is driven by non-covalent physical crosslinks, as opposed to covalent crosslinks [22]. The temperature dependence and reversibility of the solution-to-gel transition suggests that a favorable enthalpy change drives gelation, which is reinforced by the reversibility seen through re-solvation (Fig. 1C).

Next, we sought to understand the morphology and structure of CAHS D gels using scanning electron microscopy (SEM) and small-angle X-ray scattering (SAXS). SEM analysis revealed that CAHS D gels form reticular networks of interconnected fibers (Fig. 1D). This topology is similar to gels formed by gelatin, and distinct from solids formed by non-gelling proteins, such as lysozyme (Fig. 1D). SAXS analysis on the reticular mesh formed by CAHS D was performed using a concentration gradient of the protein (Fig. 1E). As a function of increasing concentration, a scattering peak emerged and increased in intensity; namely, a well-defined peak indicative of a repeating structure with a well-defined length of ∼9.5 nm (Fig. 1E, red arrow). This feature size approximates the 12.3 (+/−2.3) nm fibers observed in our SEM imaging (Fig. 1E, Fig. S1A & B), considering that preparation of materials for SEM often adds ∼2 nm of coating [23]. The large void regions between fibers are beyond the resolution of SAXS and manifest as a power law approaching small angles. Void spaces within the fibers appear as a mesh with a characteristic length scale (Fig. 1E, Fig. S1A), which shrinks in a concentration-dependent manner from ∼2.6 nm at low CAHS D concentration, to ∼2 nm at higher concentrations (Fig. S1C).

Taken together these data provide a detailed structural picture of CAHS D gels characterized by the non-covalent polymerization of CAHS D into fibers. From SAXS and SEM, we infer that a single fiber has a well-defined diameter of approximately ∼9.5 nm, but internally has less well-defined regions that lack density, the average size of which are ∼2.3 nm.

### CAHS D gels are protective and slow the motion of molecules in both the hydrated and dry state

We wondered whether gelation of CAHS D correlates with its protective capacity during desiccation. Figure 1F shows the concentration-dependent protection of the enzyme lactate dehydrogenase (LDH) [9,14] by CAHS D during desiccation (Fig. 1F). Normally desiccation and rehydration results in a loss of enzymatic activity for LDH, but when coincubated with protective molecules, LDH protection can be preserved [9,14]. While CAHS D is protective at concentrations below the point at which robust gelation is observed, optimal protection of LDH was only achieved with CAHS D at a sufficient concentration to form a continuous gel (Fig. 1F).

Given that gels can be amorphous, or ‘vitrified’ solids, we wondered if the vitrification hypothesis of desiccation could potentially help explain the protective function of CAHS D. The vitrification hypothesis, a cornerstone of the desiccation tolerance field, posits that as water is lost, protectants cause slowed molecular motion through the induction of a highly viscous state, leading to a protective, vitrified glass once desiccation is complete [24,25]. This slowing down of molecular motion and function is thought to lead to a state of reversible biostasis [24,25].

To investigate the hindrance of molecular motion in CAHS D gels, we used Fourier-transform infrared (FTIR) spectroscopy to perform hydrogen deuterium exchange (HDX) (Fig. S1D). Hydration-dependent HDX of the vitrified gel matrix was tested in dried and hydrated gels at low and high relative humidity (RH = 11% and RH = 75%, respectively; Fig. 1G, Fig. S1E). HDX experiments distinguish between a tightly packed, conformationally restricted matrix (slow exchange), and a loosely packed pliable matrix (rapid exchange) (Fig. S1D). The kinetics of the initial major phase of exchange occurred much faster in high relative humidity (minutes) than low relative humidity (tens of minutes) (Fig. 1G, Fig. S1E). The limited total exchange observed at low relative humidity indicates tight packing of CAHS D within the matrix and strong inhibition of conformational fluctuations, which decreases accessibility of peptide amide groups to the deuterated atmosphere. The higher hydration level of the matrix at high relative humidity plasticizes the gel matrix, increasing conformational fluctuations of CAHS D leading to higher amide accessibility as shown by the high percent of deuterated amide groups. When both samples were kept at RH = 100%, they achieved the same level of HDX as a solubilized protein (Fig. 1G, black dotted line), indicating that the total amides competent for exchange in each sample are equivalent when fully hydrated. These data further support the idea that molecular motion within CAHS D gels is restricted in a hydration- and concentration-dependent fashion.

The observed restricted motion of the CAHS D itself when gelled led us to ask whether a desiccation-sensitive macromolecule, such as a protein complex, could be immobilized and protected within a CAHS D gel during drying. To test this, we probed the conformational flexibility of a desiccation-sensitive photosynthetic reaction center (RC) embedded in a CAHS D gel. The RC used (purified from the photosynthetic bacterium *Rhodobacter sphaeroides* [26]) is an ideal model system to probe conformational dynamics because when an RC is immobilized, the kinetics and reaction rate distribution of the charge recombination reaction are altered (Fig. S1F) [27–29]. Conformationally restricted RCs not only have increased average rate constant (<*k*>), indicating a slowing of RC structural relaxation, but also increased width (σ) of the distribution of rate constants *k* (see eq. 1 in Methods), further indicating RC immobilization over a large range of conformational substates [27,28]. RCs embedded in vitrified CAHS D gels at RH = 11% showed increases in both <*k*> and σ (<*k*> = 35.5 s^−1^, σ = 23.8 s^−1^) compared to RC in solution (<*k*> = 8.8 s^−1^, σ = 3.6 s^−1^) (Fig. 1H, Fig. S1G & H). Increasing relative humidity in a stepwise fashion (RH = 23-53%) showed resulting stepwise increases in RC conformational mobility (Fig. 1H, Fig. S1G & H). The results observed for the conformational dynamics of RCs incorporated into vitrified CAHS D matrices were similar to those obtained with RCs embedded into glassy matrices formed by trehalose (Fig. S1G & H), a sugar known to function as an extraordinary bioprotectant via immobilization of sensitive biological material in a vitrified state [28,30]. Interestingly, CAHS D and trehalose glassy matrices incorporating RCs exhibit very similar water absorption properties (Fig. S1I).

Combined, these data demonstrate that optimal protection during drying is only achieved at concentrations where CAHS D gels, and that gelation restricts the conformational flexibility of CAHS D itself as well as the molecular motion of larger desiccation-sensitive macromolecules embedded within the gel-matrix. These findings lead us to hypothesize that gelation may play a mechanistic role in tardigrade desiccation tolerance through the immobilization of sensitive biomolecules, which could induce biostasis.

### Characterization of the CAHS D global structural ensemble

To begin to understand the mechanisms underlying CAHS D gel formation, which would allow us to experimentally probe the role of gelation in stress tolerance, we performed all-atom Monte Carlo simulations of monomeric CAHS D. The simulations are consistent with a largely disordered protein, and suggest that CAHS D occupies a dumbbell-like conformational ensemble, in which the *N-* and *C-*termini of the protein are relatively collapsed and are held apart by an extended linker region (Fig. 2A).

**Figure 2.**
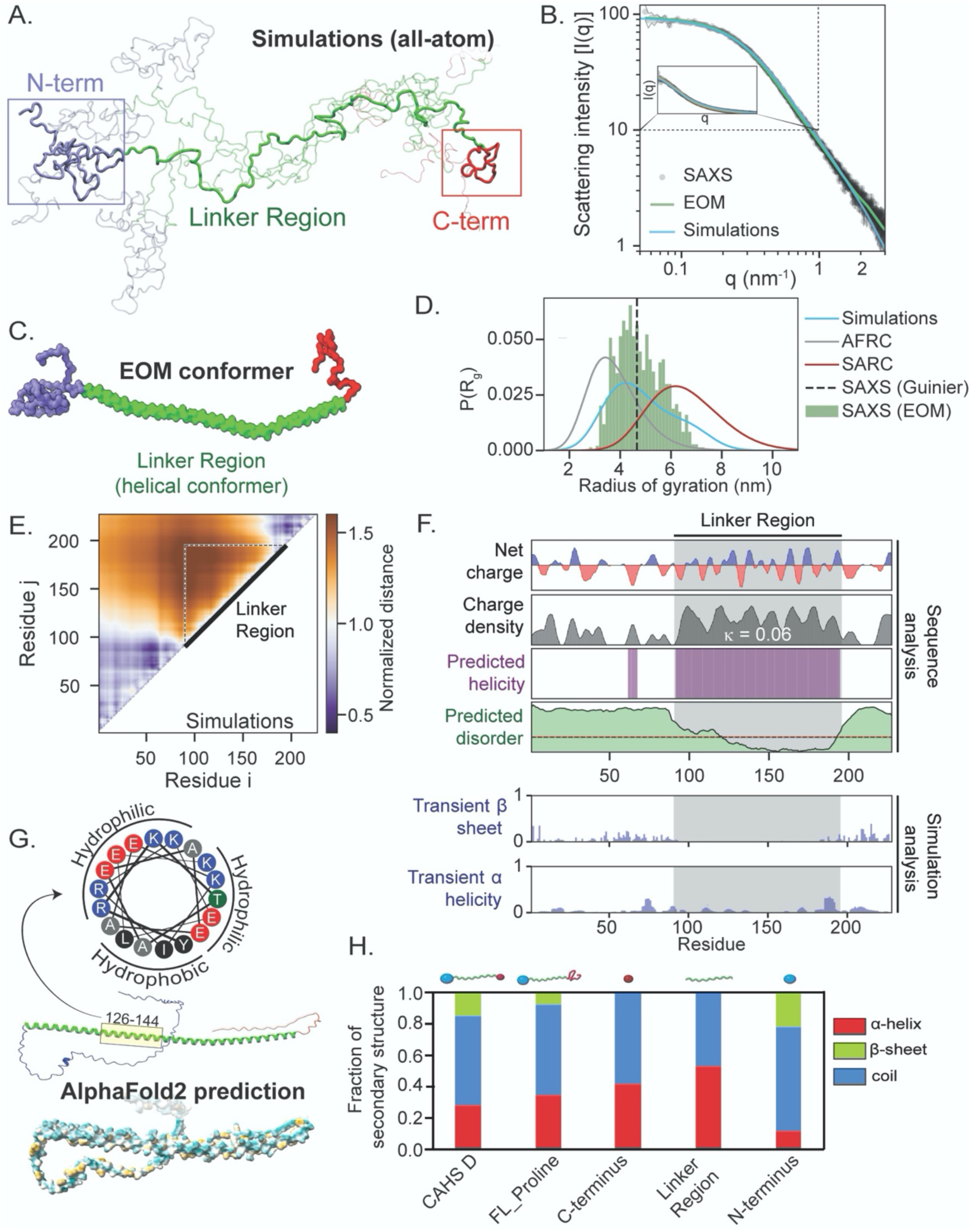
Global and local ensemble features of CAHS D. **A)** Representative depiction of the CAHS D ensemble derived from all-atom Monte Carlo simulation. Simulations predicts two collapsed termini held apart by an extended linker. **B)** SAXS data (black), EOM fitted scattering profile (green), and simulation-derived scattering profile (blue) superimposed with one another. EOM and simulations are in good agreement with experimental data. See also Fig. S2B-F. **C)** Representative depiction of the CAHS D ensemble derived from EOM analysis. The three domains observed in simulations (A) are highlighted in blue (N-terminus), green (linker), and red (C-terminus). **D)** Comparison of the radius of gyration distributions for monomeric CAHS D obtained from SAXS (dotted line), EOM (green bars), and Monte Carlo simulation (blue curve). For comparison, overlays for the Gaussian-chain-like AFRC model (grey) and self-avoiding random coil (red) generated from sequence-matched ensembles are shown. **E)** A normalized distance map from CAHS D simulations compares the average inter-residue distance with the expected inter-residue distance if CAHS D behaved like a Gaussian chain. Hotter colors are further apart, cooler colors are closer. This analysis suggests that the terminal regions are largely collapsed, while the linker is expanded. **F)** Localization of different sequence features within the CAHS D ensemble using bioinformatics and simulation analysis. **G)** (bottom) Alphafold2 predictions of the CAHS D ensemble suggest high amphipathic helical propensity within the linker region (dark cyan most hydrophilic, dark golden most lipophilic). Helical wheel diagram of residues 126-144 of CAHS D (top). **H)** Circular dichroism spectroscopy of CAHS D showing the fraction of different types of secondary structure for different variants.

To assess predictions of the global ensemble of CAHS D, we performed size exclusion chromatography-coupled SAXS (SEC-SAXS) of monomeric CAHS D. The linearity of the low-q region after Guinier transformation supports the monodispersity of this sample (Fig. S2A). The ensemble radius of gyration from the Guinier approximation is 4.66 ± 0.07 nm. The scattering data were also analyzed using the Ensemble Optimization Method (EOM) to generate a SAXS-derived distribution of radius of gyration (R_g_) values. From the EOM ensemble, the average R_g_ is 4.84 ± 0.92 nm, which is slightly larger than that estimated from the Guinier analysis (Fig. 2B, Fig. S2B-E). Consistent with these experimental results, the R_g_ predicted by ALBATROSS (a deep-learning based sequence-to-ensemble predictor) is 4.75 nm [31]. Furthermore, EOM analysis suggests that CAHS D can sample dumbbell-like conformations, in good agreement with the ensemble generated from Monte Carlo simulations (Fig. 2C & D, Fig. S2E).

Next, to improve the rigor of the CAHS D structural model, we refined the simulation ensemble using the experimental SAXS data. We employed a Bayesian/Maximum-Entropy (BME) reweighting protocol in which the simulation frames are weighted such that the observables from the new simulation ensemble match those from the ensemble-averaged SAXS data [32]. Unless otherwise specified, all ensuing simulation analyses used the subsampled and reweighted ensemble (Figs. S2F & G) (see Materials and Methods and SI).

The CAHS D ensemble dimensions are highly similar between simulations and experiments (Fig. 2D). Notably, CAHS D is more compact than a self-avoiding random coil (SARC Rg, = 6.5 nm) and more expanded than a sequence-specific Gaussian chain model (the analytical Flory random coil, AFRC) of the same length (AFRC R_g_ = 3.8 nm) (Fig. 2D) [33]. We compared the inter-residue distances in the CAHS D simulations to those from the AFRC model and determined regions of local expansion and compaction. From the normalized distance map, the *N-* and *C*-termini are, on average, more compact than in the AFRC model, whereas the linker (residues 91 – 195) is more expanded (Fig. 2E). Considering also the global CAHS D dimensions, these data implicate the Linker Region as the overall driver of ensemble expansion.

Analysis of the CAHS D sequence supports a linker charge-driven model to spatially separate the *N-* and *C*-termini. The sequence parameter kappa (κ) ranges from zero to one and quantifies the charge patterning in a protein sequence, such that linearly well-mixed sequences have low κ values, and sequences with segregated blocks of like charges have high κ values [34,35]. Full-length CAHS D is highly charged, and those charges are well-mixed (κ = 0.086). However, the Linker Region has a lower κ of 0.06, making it more well-mixed than over 99% of all tardigrade IDRs (Fig. S2H&I). In other systems, highly charged well-mixed IDRs (low κ) are more expanded than those within which charged residues are segregated (high κ) [34–36]. Taken together, our simulations, SAXS, and bioinformatics analyses suggest that CAHS D samples a dumbell-like ensemble composed of two collapsed terminal regions bridged by a highly extended Linker Region, preventing intramolecular interaction between the *N-* and *C*-termini, and poising CAHS D for intermolecular interaction.

### Characterization of the CAHS D local structural ensemble

While our simulations predict CAHS D is disordered, this does not preclude the existence of transient structure. Our simulations suggest low levels of transient β-strand in the *N*- and *C*-terminal regions, while the Linker Region is predicted to contain transient α-helix (Fig. 2F, Fig. S2J). Our simulations likely underestimate this helicity – bioinformatic predictions strongly indicate that the Linker Region is substantially helical, which is supported by recent solution-phase NMR measurements that reveal transient helicity (Fig. 2F, Fig. S2J) [19].

Furthermore, the Linker Region is predicted to be an amphipathic α-helix with hydrophilic and hydrophobic faces (Fig. 2G). We probed this structure experimentally in solution for each region of CAHS D using circular dichroism (CD) spectroscopy (Fig. 2H, Fig. S2K). CD spectra at non-gelling concentrations confirm the largely disordered nature of full-length CAHS D, with some propensity for A-helix and β-structure formation (Fig. 2H). However, at concentrations when gelation is observed, CAHS D showed increased α-helical character with the helix fraction ∼44% at 700 μM (17.7 mg/mL) which is close to the fraction of 46% expected for CAHS D with a fully α-helical Linker Region (Fig. 1B, Fig. S3E). Consistent with bioinformatic predictions, CD spectroscopy of truncation mutants of each CAHS D region confirmed substantial helical content in the Linker Region (∼40%) (Fig. 2H). Similarly to CAHS D, the Linker Region protein gave a strong α-helical signal that increased at high concentrations to 89% helix at 700 μM (8.6 mg/mL) (Fig. 3C, left panel, Fig. S3F). Such high helix fractions for both CAHS D and the Linker Region at increased concentrations suggest that self-assembly of CAHS D resulting in gelation is accompanied by the formation of helical bundles of the Linker Regions.

**Figure 3.**
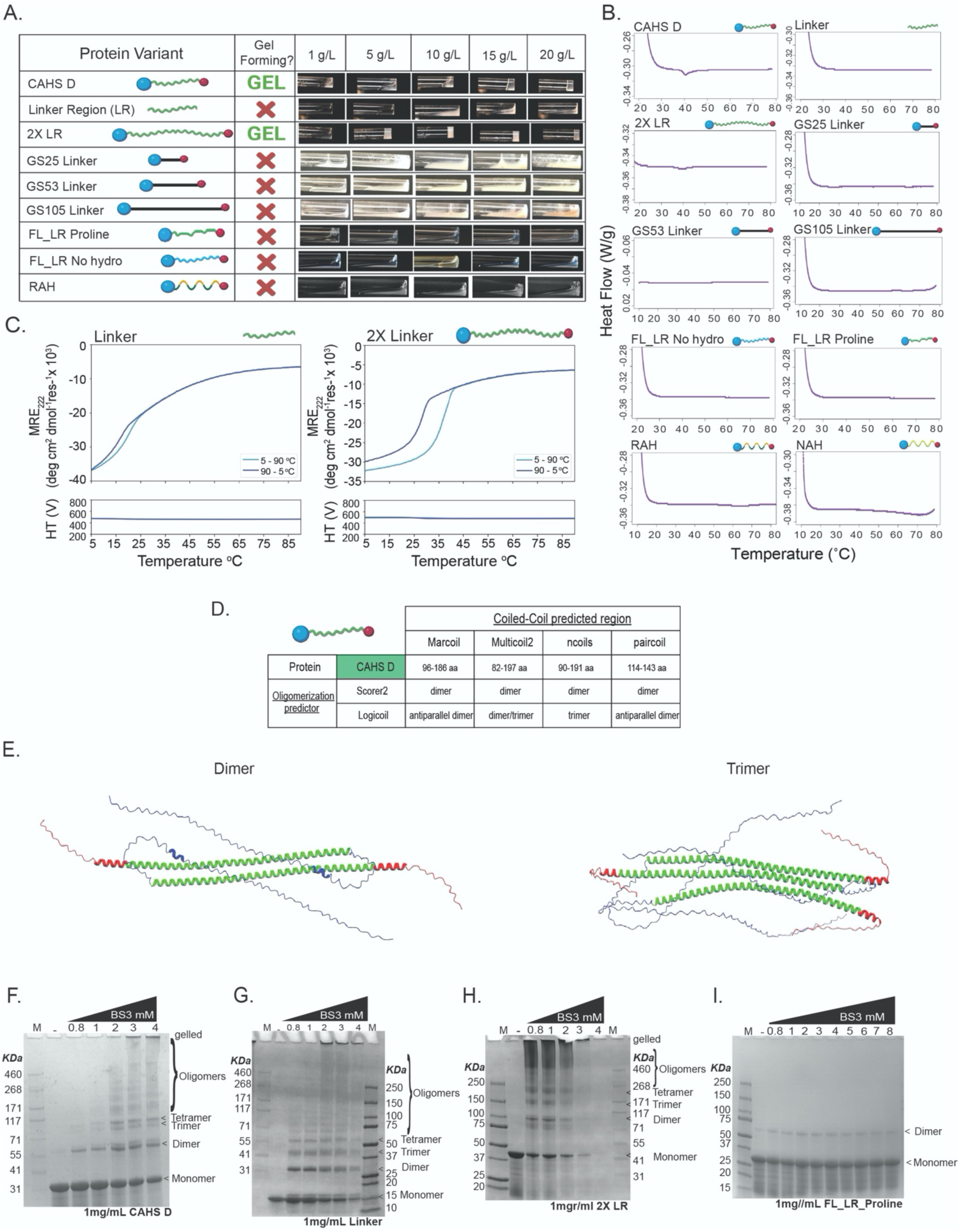
CAHS D’s Linker Region is necessary but not sufficient for gel formation. **A)** Table with graphical representations of CAHS D variants (column 1), whether or not they form a gel under the conditions tested (column 2), and images of solutions/gels formed by the variants at various concentrations (column 3). **B)** Differential scanning calorimetry melt curves for CAHS D and its linker variants at 20 mg/mL. Only CAHS D and 2X Linker show melts characteristic of gels. **C)** Thermal responses of the CD signals at 222 nm (ramping up, light blue line; and ramping down, dark blue line) for Linker Region (0.7 mM, 8.6 mg/mL) and 2X Linker Region (0.25 mM, 9.3 mg/mL) variants. Note that in both cases self-association is observed, revealing that while the Linker Region alone can self-associate it cannot form a gel. **D)** Coiled-coil predictions for CAHS D from a variety of predictors. **E)** AlphaFold-Multimer prediction of double and triple CAHS D complexes showing antiparallel arrangement of the molecules. **F)** SDS-page gel of 1 mg/mL CAHS D incubated with increasing amounts of BS3 amine ester crosslinker to visualize oligomeric state of CAHS D. G) SDS-page gel of 1 mg/mL Linker Region incubated with increasing amounts of BS3 amine ester crosslinker to visualize oligomeric state of Linker Region. H) SDS-page gel of 1 mg/mL 2X Linker incubated with increasing amounts of BS3 amine ester crosslinker to visualize oligomeric state of 2X Linker. I) SDS-page gel of 1 mg/mL FL_LR_Proline incubated with increasing amounts of BS3 amine ester crosslinker to visualize oligomeric state of FL_LR_Proline. In all crosslinking gels, the first lane has MW marker, the second lane has 1 mg/mL of proteins with no crosslinker showing monomeric state. Lanes 3 to end have 1 mg/mL of crosslinked proteins with increasing amounts of BS3 (0.8 mM to 4 mM / 8 mM) showing formation of dimers, trimers and high order oligomers.

Consistent with secondary structural prediction, CD spectroscopy indicates some β-structure in the truncated *N*-terminal Region (Fig. 2H). Contrary to the secondary structure predictions, we did not observe residual β-structure by CD spectroscopy in the isolated *C*-terminal Region. However, we reasoned that the low β-structure content observed by CD spectroscopy in the isolated *C*-terminus could be caused by the loss of sequence context altering the conformational ensemble [37–40]. To test this possibility, we generated an additional CAHS D variant (FL_Proline). The FL_Proline mutant was designed to disrupt secondary structure in the *C-*terminus via the insertion of three prolines into the three predicted β-strands (Fig. S2J) [41,42]. The CD spectrum of FL_Proline showed >50% reduction in β-structure content relative to wildtype (Fig. 2H, S2L), suggesting that the *C* terminus does have β-structure when in the context of the full-length CAHS D protein. Moreover, we found FL_Proline to be defective in gel formation (Fig. S2M).

Taken together, the results from CD spectroscopy support computational analysis and show helical content throughout CAHS D, with some localized, but likely transient, β-structure content in the terminal regions.

### The amphipathic nature of the Linker Region drives helical interactions and bundling of CAHS D

With a general understanding of both the global and local ensemble properties of CAHS D, we set out to understand i.) which regions of CAHS D are necessary and sufficient for gel formation, and ii.) which secondary structural elements help mediate this process.

We began by testing the necessity and sufficiency of the Linker Region and its properties in mediating gelation. To this end, we generated a Linker Region (LR) variant lacking both the *N*- and *C*-terminal regions (Fig. 3A). This variant failed to form gels (Fig. 3A & B), demonstrating that the Linker Region alone is not sufficient for gelation.

Next, we generated a variant we term 2X Linker Region (2X LR), which has the wildtype *N*- and *C*-termini held apart by a tandem duplication of the wildtype Linker Region (Fig. 3A). 2X LR gelled at a lower concentration than the wildtype CAHS D (Fig. 3A & B) suggesting that while the Linker Region is not sufficient for gelation, it does play a role in the process.

Since gelation at a lower concentration with our 2X LR variant could be due to additional interactions between the extended Linker *or* to increased distance between terminal regions, we generated a series of glycine-serine (GS) repeat variants. These GS variants replaced the endogenous Linker Region with GS repeats resulting in Linkers with predicted smaller (GS25; R_g_ = 4.04 nm), equivalent (GS53; R_g_ = 4.65 nm), or larger (GS105; R_g_ = 5.61 nm) R_g_ than the wildtype Linker Region (Fig. 3A) [43,44]. None of these variants gelled (Fig. 3A & B) showing that sequence features and properties of the endogenous Linker Region, beyond merely holding terminal regions apart, are necessary for gelation.

To assess which sequence features of the Linker Region are essential for gelation, we began by testing the necessity of the linker’s helicity in mediating gelation. Proline residues were inserted every 6 – 8 amino acids within the Linker Region to break the helicity of the full-length CAHS D (FL_LR Proline; Fig. 3A). As expected, we observed that these mutations led to a decrease in helicity (Fig. S3A) and our FL_LR Proline variant failed to gel (Fig. 3A & B), again indicating helicity of the Linker Region is important for this process.

We probed the characteristics of the wildtype Linker Region helix that are important for gelation further by assessing the sequence chemistry of the Linker Region in mediating this process. To this end, we generated a variant consisting of the full-length protein with all the hydrophobic residues in its Linker Region substituted for glycine residues (FL_LR No hydro), which also failed to gel (Fig. 3A & B, Fig. S3B). Next, we generated two variants that maintained the hydrophobicity of the Linker Region, while shuffling the distribution of hydrophobic and hydrophilic residues resulting in a non-amphipathic helix (NAH) or a robust amphipathic helix (RAH), both of which also failed to gel (Fig. 3A & B, Fig. S3C & D). We note that the NAH variant expressed and purified with extremely low yield, leading us to characterize its gelation only by DSC, due to the large input of material required for direct observation as presented in Figure 3A.

Taken together, these results demonstrate that while the Linker Region is not sufficient for gelation, it is necessary, and furthermore that the predicted amphipathic helix within this region is highly tuned to mediate this phenomenon.

We speculated that the Linker Regions of different CAHS D proteins could interact with one another to help mediate gelation. As mentioned above, the high helix fractions for both CAHS D (44%) and LR (89%) at increased concentrations suggest that self-assembly of CAHS D resulting in gelation might be accompanied by the formation of helical bundles of the Linker Region. To test this, we performed CD spectroscopy on CAHS D and both the LR and 2X LR variants at concentrations at which wildtype CAHS D forms robust gels in controlled heating and cooling experiments (Fig. 1B and Fig 3C). All three proteins showed cooperative unfolding upon heating indicative of self-assembly (Fig. 1B and Fig 3C). Both processes were highly reversible (Fig. S3E-G). Interestingly, the midpoint melting temperature (T_M_) of 2X LR variant was similar to that for CAHS D (36 – 37 °C); *i.e.,* linker length does not affect the melting point (Fig. 1B & Fig. 3C). The LR alone at high concentrations showed a similar cooperative transition, but with a significantly lower T_M_ of 20 °C. This confirms the significant role of the terminal domains for the self-assembly and gelation of CAHS D and 2X LR proteins.

Next, we wondered what the nature of the Linker-Linker interaction might be. Since CAHS proteins possess amphipathic helices with strong hydrophilic and hydrophobic faces (Fig. 2G) [15,17], and since the burial of hydrophobic residues is known to be a strong driver of A-helical coiled-coil formation [45,46], we assessed the predicted ability of CAHS D to form coiled-coils using four different prediction tools for coiled-coil propensity and oligomerization (Fig. 3D, Fig. S3H). CAHS D is predicted to form coiled-coil dimers and trimers, with antiparallel dimers being predicted by most tools (Fig. 3D). Furthermore, AlphaFold-Multimer predictions with two or three copies of the CAHS D sequence show antiparallel helical interactions between Linker Regions (Fig. 3E).

While predictive tools largely suggest an antiparallel dimeric assembly of CAHS D, we directly assessed the oligomerization state of CAHS D using an amine ester crosslinker (Fig. 3F). SDS-PAGE denaturing gels show a formation of dimers at low crosslinker concentration and higher order oligomers (including faint bands indicating trimerization) upon an increase of crosslinker concentration.

We also assessed the oligomerization state of the Linker Region (Fig. 3G), 2X LR (Fig. 3H) and FL_LR_Proline (Fig. 3I) by amine ester crosslinking. The Linker Region shows formation of dimers, trimers and tetramers at low concentration and higher order oligomers upon increasing crosslinker concentration. This means the Linker Region by itself is enough to drive helix-helix interactions. 2X LR shows even at low crosslinker concentration of 0.8 mM, dimers and higher-order oligomers, in line with the capacity of this protein to oligomerize and form gels at low concentrations. FL_LR_Proline variant shows very faint dimer bands, but even without the crosslinker. Breaking the helicity in the Linker region of CAHS D disrupts oligomerization of the protein

Combining data from the experiments described above, we conclude that the Linker Region of CAHS D is essential for its gelation and that gelation, at least in part, relies on the helicity and amphipathic nature of the linker to form antiparallel dimers.

### β-structure Interactions between CAHS D termini drive gel-network formation

Since the Linker Region of CAHS D is necessary but not sufficient for gelation, we wondered which other region(s) of the protein are required to drive gel formation.

To test the necessity of the terminal regions of CAHS D in mediating gelation, we produced variants that have either deleted or swapped N- and/or C-terminal Regions (Fig. 4A). We observed that removal of either the *N*- or the *C*-terminal Region (NL1 and CL1), or disruption of the *C-*terminus by inserting proline residues (FL_Proline variant) resulted in a failure to gel (Fig. 4A & B, Fig. S2M, Fig. S4A). Likewise, replacing the *N-*terminal Region with the *C*-terminal Region (or *vice versa,* NLN and CLC variants) resulted in a loss of gelation (Fig. 4A & B, Fig. S4A), implying that heterotypic interactions between the two regions are likely essential for gelation, in line with our observation that helical interactions within the Linker Region are predicted to take place in an antiparallel fashion (Fig. 3D & E).

**Figure 4.**
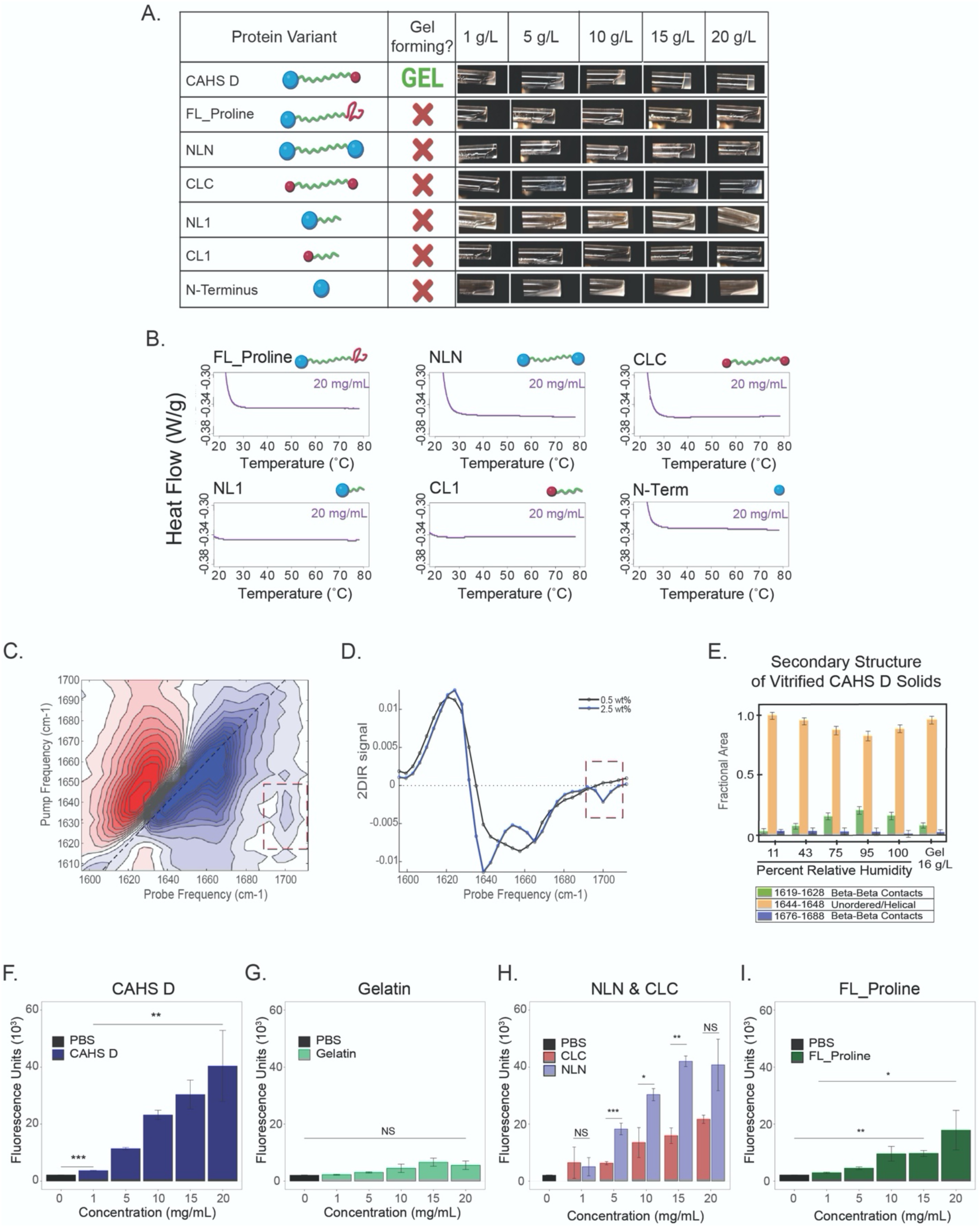
Heterotypic terminal domains are required for CAHS D gel formation. **A)** Table with graphical representations of CAHS D variants (column 1), whether or not they form a gel under the conditions tested (column 2), and images of solutions/gels formed by the variants at various concentrations (column 3). **B)** Differential scanning calorimetry melt curves for CAHS D and its terminal variants at 20 mg/mL. **C)** 2D-IR spectrum of CAHS D at a concentration above the gelation threshold (2.5 wt%). **D)** horizontal slice through the 2D-IR spectrum (obtained by averaging over the pump-frequency range from 1625 to 1635 cm^−1^) of 0.5 wt% (black line) and 2.5 wt% (blue line) of CAHS D. Red dotted square shows the emergence of β-sheet at 2.5 wt%. **E)** Relative content in secondary structures of the CAHS protein as determined by FTIR analysis of the amide I’ band in a glassy matrix at different hydration levels (equilibrated at a relative humidity from 11% to 100%, as indicated) and in the hydrated gel state at concentrations of 16 g/L. The Gaussian sub-bands centered in the wavenumber interval (1644-1648 cm^−1^) are attributed to unordered regions, while the Gaussian components peaking in the wavenumber intervals (1619-1628 cm^−1^) and (1676-1688 cm^−1^) are indicative of interprotein β-sheet structures. **F)** Thioflavin T (ThT) fluorescence as a function of concentration for CAHS D, **G)** gelatin, **H**) NLN, CLC and **I)** FL_Proline. Error bars represent +/−1 standard deviation. Significance determined using a Welch’s t-test. All experiments presented used a minimum of 3 replicates. Asterisks represent significance relative to wild type CAHS D or buffer controls as indicated. *p<0.05, **p<0.01, ***p<0.001, ****p<0.0001, NS is not significant.

To determine what ensemble structure within the termini drives gel formation, we performed femtosecond two-dimensional infrared (2D-IR) spectroscopy on CAHS D solutions at concentrations below and above the critical concentration for gelation. 2D-IR cross peaks are reliable markers for the presence of β-structure [47,48], especially in the presence of other secondary structures in the protein [49]. Our analysis shows the emergence of β-structure in gelled, but not non-gelled CAHS D (Fig. 4C & D).

Next, we assessed how the conformation of CAHS D changes going from the gelled to the desiccated state using FTIR. We observed that β-structure increased in the hydrated state as a function of concentration (Fig. 4D). This increase continued in the drying hydrogel (95 - 100% RH) but decreased at lower hydration levels (11 – 75% RH) (Fig. 4E). This implies that there is an optimal hydration level for stabilizing β-structured contacts, which may relate to the need for higher stability while the matrix is undergoing the final stages of drying or the early stages of rehydration.

Finally, to determine whether higher β-β interactions correspond with the expected β-structure content of the termini, and the ability of these to interact, we assayed CAHS D at different concentrations with thioflavin T (ThT). ThT is used as a fluorescent indicator of amyloid fibrils [50,51], but more generally reports on β-structured assemblies [52–54]. We observed steady increases in ThT fluorescence intensity as a function of CAHS D concentration (Fig. 4F), consistent with higher β-structure as gelation progresses. ThT signal in samples containing gelatin, which forms gels exclusively via entwined ABhelices without the need for β-β contacts [55], was not significantly different from buffer controls (Fig. 4G).

To determine how each terminus contributes to β-β interactions, we probed three variants using the ThT assay; NLN and CLC, which have two homotypic termini, and FL_Proline, which has disrupted *C*-terminal β-structure (Fig. 4A, Fig. 2H, Fig. S2L). ThT labeling of NLN and CLC variants showed a concentration-dependent increase in fluorescence, with higher fluorescence observed in NLN than CLC (Fig. 4H), which is consistent with the predicted and measured degree of β structure in each terminus (*N versus C*) (Fig. 2H). The Tht fluorescence of FL_Proline was less than variants with two endogenous termini but more than the gelatin control, as would be expected of a protein that is only competent to form β-β interactions with one terminus (Fig. 4I). It is important to note that ThT fluorescence does not report on the strength of interactions, simply the relative quantity of interactions. Therefore, the pattern observed for ThT fluorescence; NLN>CAHS D>CLC>FL_Proline, follows the predicted β-structure content of these variants (Fig. 2H). Combined with our determination that NLN and CLC variants do not gel, these data suggest that while N-N and C-C interactions can occur they are not able to induce gelation as is observed with N-C interactions.

Taken together, these data suggest that heterotypic interactions between β-structure in the *N*- and *C*-terminal Regions of CAHS D are necessary for the gel-forming abilities of this protein. However, neither the *N-* or *C-*termini are sufficient themselves for gelation (Fig. 4A & B).

### Gelation of CAHS D occurs through helical interactions of the linker and end-to-end polymerization via β-β interactions

Based on the experiments and data described above, we conclude that both the interactions between Linker Regions and β-β interactions between heterotypic terminal regions of CAHS D are necessary to form dimers that ultimately assemble into higher-order structures, *i.e.*, fibers with a diameter of ∼9.5 nm.

It has been proposed that gelation of CAHS proteins might be mediated exclusively by the stacking of helices [19] or coiled-coil interactions between the *C*-terminal ends of these helices [20] (Fig. 5A). Our empirical demonstration that CAHS D gelation requires both its *N*- and *C*-terminal regions rules out these possibilities as the exclusive mechanisms mediating CAHS D gelation. Nevertheless, we probed the mechanism of fiber formation further by examining the morphology and dimensions of gels formed by our 2X Linker variant.

**Figure 5.**
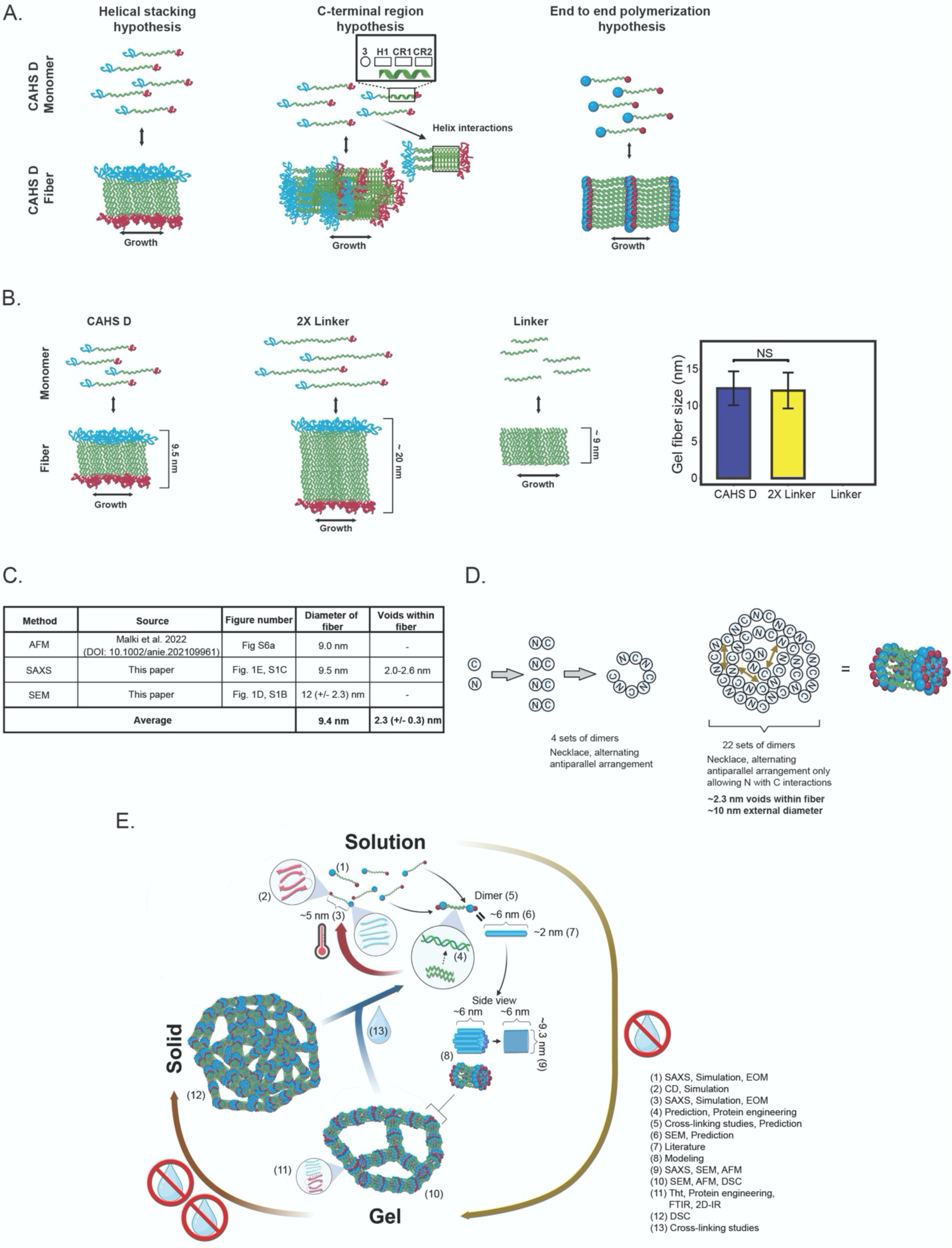
Model of *in vitro* CAHS D gel formation. **A)** Schematic representation of leading hypotheses on how CAHS gel formation could be mediated. **B)** Schematic representation of expected gel fiber diameters given the helical stacking hypothesis (left). Quantification of fiber diameters from CAHS D and 2X Linker gels (right). **C)** Table summarizing empirical measurements of CAHS D gel fiber diameter. **D)** Modeling of a necklace arrangement of CAHS D monomers and dimers. **E)** Working model of CAHS D dimerization and gel formation. Numerical annotations note where evidence for model elements come from.

In the “helical stacking” hypothesis, one would expect that the width of gel fibers should be approximately doubled in our 2X Linker variant as compared to the wildtype variant (Fig. 5B). However, quantitative analysis of scanning electron micrographs of fibers formed by wildtype CAHS D and the 2X Linker variant showed no statistical difference in gel-fiber width (12.3 +/−2.3 *versus* 12.1 +/−2.4 nm, respectively) (Fig. 5B, Fig. S5A & B). One would also expect that if the termini do not play a role in gelation as stated in the helical stacking hypothesis, our Linker Region variant would also gel (Fig. 5B), but it does not (Fig 3A & B). Additionally, simulation, SAXS, and EOM indicate that the R_g_ of monomeric CAHS D is ∼5 nm, well short of the ∼9.5 nm fiber diameter as measured by SAXS (Fig. 1E), SEM (Fig. S1B), and AFM [19]. This logic would suggest that the helical stacking hypothesis is unlikely, since it does not fit with the requirement for the terminal regions, nor is it congruent with empirically determined dimensions of CAHS D gel fibers. Similarly, a mechanism by which CAHS D gels exclusively through helical interactions in the *C* terminus can be ruled out due to the necessity of having both the *N*- and *C*-terminal regions of the protein as well as the necessity of β-structured interactions between these regions (Fig. 4A-I).

Combining results from global and local ensemble studies, protein engineering, imaging, material and biophysical characterization, we arrive at a model of CAHS D gelation that is consistent with empirical measurements of CAHS D fiber measurements (Fig. 5C-E). In solution, monomeric CAHS D forms antiparallel, staggered, dimers with a length of ∼6nm (Fig. S5A & B) and other higher-order structures in a concentration-dependent fashion through the association of linker regions of adjacent CAHS D proteins (Fig. 3D, 3F & Fig. S3H). Considering the ∼9.4 nm diameter and ∼2.3 nm voids of gel fibers (Fig. 1E, Fig. S1C), we invoke a necklace model (Fig. 5D) in which 22 dimers associate and supercoil into a bundle with these dimensions. Bundles of antiparallel dimers then polymerize through end-to-end β-β interactions to form a gel (Fig. 5E).

Further desiccation of the gel results in solidification into a non-crystalline amorphous solid [9,56], which is expected as gels themselves are amorphous solids [57,58]. Finally, rehydration results in a rapid re-solvation and dissipation of the gel as CAHS D molecules rapidly go back into solution (Fig. 5E, Fig. S5C & D).

### CAHS D forms gel-like condensates in living cells during osmotic stress

CAHS proteins from a different tardigrade species have been reported to form condensates in cells upon osmotic shock, and condensate formation stiffens cells [20]. However, the sequence features underlying this condensation have not been fully elucidated nor has the role of condensation in cellular stress been tested empirically [20,21]. To assess whether the ability of CAHS D to condense into a gel *in vitro* is reflected by its *in vivo* behavior, and to understand the sequence features that drive this behavior, we generated a stable Human embryonic kidney (HEK) cell line expressing an mVenus:CAHS D fusion protein. We also generated a stable cell line expressing only mVenus protein as control.

Both cell lines expressing mVenus alone and our mVenus:CAHS D fusion show diffuse distributions of protein in the cytoplasm of non-stressed HEK cells (Fig. 6A). However, while the localization of mVenus remained diffuse within the cytoplasm of osmotically shocked HEK cells, mVenus:CAHS D localized into fiber-like condensates (Fig. 6A).

**Figure 6.**
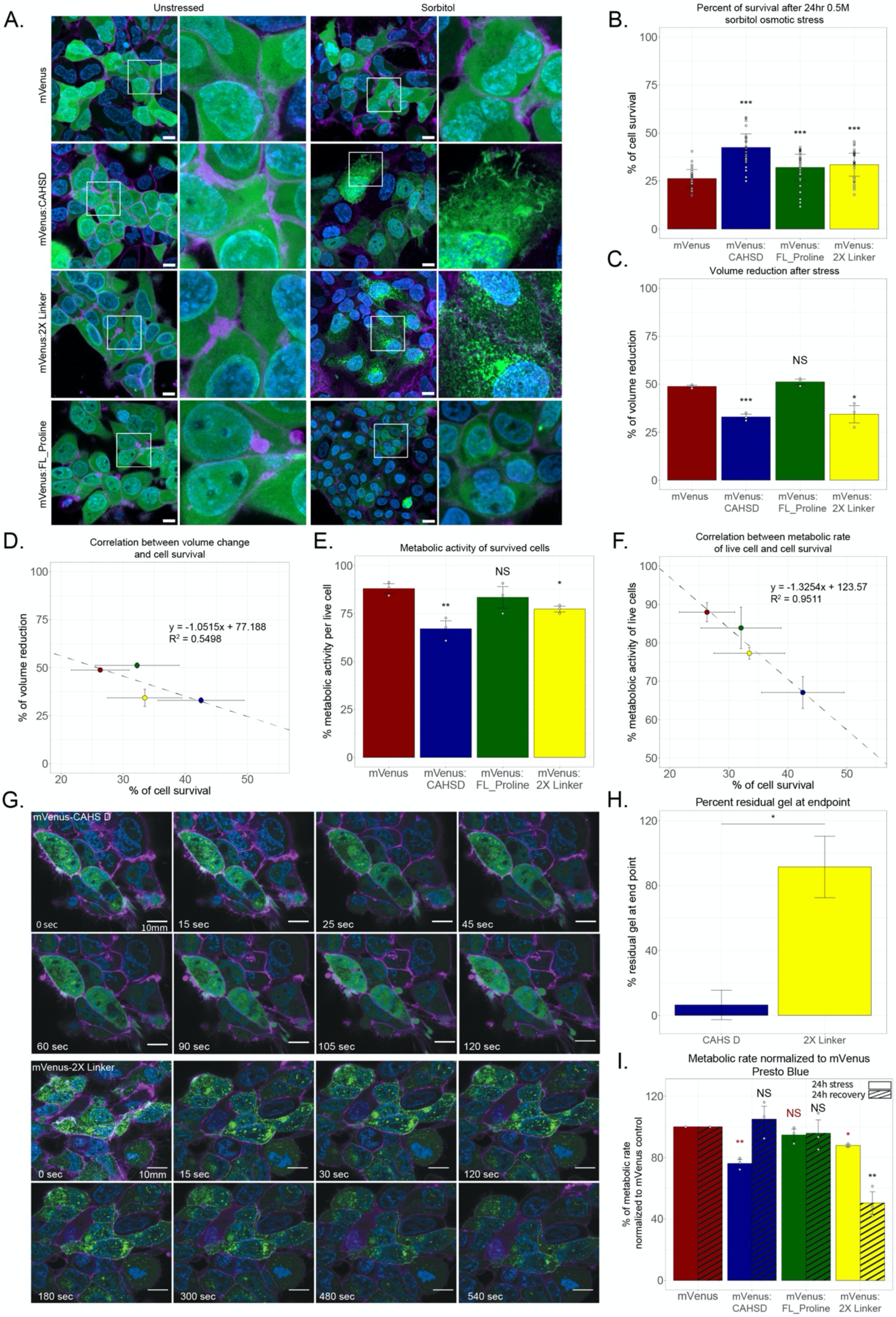
Condensation of CAHS D induces reversible biostasis and increases survival during osmotic shock. **A)** Fluorescence microscopy of human embryonic kidney cell lines expressing mVenus or mVenus fusions to CAHS D variants. The left two columns show cells and insets of cells imaged under non-stressed conditions. The right two columns show cells and insets of cells treated with 0.5 M sorbitol. Green = mVenus, blue = DNA, purple = cell membrane. Scale bar = 10 µm. **B)** Combined survival data from Calcein AM/Propidium iodide and Hoechst/Propidium iodide survival assays (data from each individual assay shown in Fig. S6C & D). **C)** Cell volume reduction during osmotic stress. **D)** Correlation between cell volume change and survival during osmotic stress. **E)** Quantification of metabolic activity of living cells during osmotic stress using Presto Blue HS assay (For XTT metabolic assay data see Fig. S6I & J). **F)** Correlation between reduction in metabolic activity and survival during osmotic stress. **G)** Confocal time-lapse imaging of CAHS D and 2X Linker condensates in cells recovering from osmotic stress. Scale bar = 10 µm. **H)** Quantification of residual condensational at end-point of time-lapse experiment (120 seconds for CAHS D and 540 seconds for 2X Linker). **I)** Normalized metabolic rates of alive cells to mVenus control cells during osmotic stress (solid bars) and after 24h recovery (striped bars) (For XTT assay data see Fig. S6J). Error bars represent average deviation. Significance determined using a paired T. Test, asterisks represent significance relative to mVenus control cells. *p<0.05, **p<0.01, ***p<0.005, NS is not significant. In figure I red color statistics represent significance relative to mVenus stressed cells, and black color statistics represent significance to mVenus recovered cells.

Next, we assessed if sequence features that promote gelation of CAHS D *in vitro* also drive condensation *in vivo*. For this, we established stable HEK cell lines expressing fusions of mVenus and the FL_Proline and 2X Linker variants and observed their subcellular localization before and during osmotic stress. Both mVenus:FL_Proline and mVenus:2X Linker were diffusely distributed through the cytoplasm, similar to mVenus and mVenus:CAHS D, in unstressed cells (Fig. 6A, Fig. S6A & B). However, upon osmotic shock mVenus:2X Linker assembled into fibrillar condensates, while mVenus:FL_Proline remained diffuse (Fig. 6A, Fig. S6A & B).

This pattern of condensation matches our *in vitro* observations, where disruption of terminal regions results in a loss of gelation, while tandem duplication of the linker region results in gelation at even lower concentrations than the wildtype protein (Fig. 3A & B). Quantification of condensate formation showed robust statistical increases in fiber formation only in CAHS D variants that form gels *in vitro* (Fig. S6A & B). These data indicate that the same sequence features driving *in vitro* gelation of CAHS D are responsible for its *in vivo* condensation during osmotic stress.

### CAHS D promotes survival during osmotic shock in HEK cells

Since the gelled state of CAHS D has been shown to modulate its protective capacity under different types of desiccation stress [18,59], we wondered if gelation of CAHS D in cells during osmotic shock might provide some degree of increased protection. To assess this, we utilized two different cellular survival assays on osmotically stressed cells expressing our mVenus, our wildtype CAHS D protein, and our gelling (2X Linker) and non-gelling (FL_Proline) variants (Fig. S6C & D).

After 24 hours of 0.5 M sorbitol osmotic shock, we observed a large decrease in cell viability for mVenus-expressing cells (Fig. 6B, Fig. S6C-E). Cells expressing FL_Proline or 2X Linker survived significantly better than mVenus alone (Fig. 6B). However, CAHS D retained the highest viability, which was significantly higher than cells expressing mVenus as well as our gelling and non-gelling variants of CAHS D (2X Linker and FL_Proline).

To ensure that the increase in survival seen in osmotically shocked cells expressing CAHS D and its variants is not due a cellular response from an induced ER Stress or unfolded protein response (UPR), we measured the level of ER stress/UPR on the cell lines using a ratiometric stress sensor [60]. Expression of mVenus:CAHS D or any of its variants does not induce significant levels of ER stress/UPR compared to naive HEK cells or HEK cells expressing mVenus alone (Fig S6F).

While wildtype CAHS D was significantly more protective than either of its variants, the observation that all CAHS D variants used in this study resulted in increased cellular survival during osmotic shock, regardless of their ability to form gels, suggests that CAHS proteins are protective in both their condensed and uncondensed state *in vivo*.

### Condensed CAHS D serves as a stress-induced cytoskeleton to preserve cellular ultrastructure

The observation that regardless of their condensed state CAHS proteins are protective in cells during osmotic shock intrigued us. This observation is in line with reports that *in vitro,* CAHS D provides protection in both its soluble and gelled form albeit *via* distinct mechanisms [18,59].

To probe what protective mechanisms might be conferred by condensed CAHS D in cells during osmotic shock, we examined the role of CAHS D condensates in helping maintain cell volume during osmotic shock induced cell shrinkage. This investigation was prompted by the observation that the mechanism by which CAHS D and other CAHS proteins form gels is in some respects reminiscent of proposed mechanisms underlying Intermediate Filament (IF) formation [61–64]. As such, we wondered whether CAHS D might function as a novel type of stress-inducible cytoskeletal protein, helping to ameliorate the dramatic changes in cell volume associated with hyperosmotic shock.

To assess whether condensation of CAHS proteins can help preserve the structural integrity of cells, we examined volumetric changes associated with osmotic shock in cells expressing gelling and non-gelling versions of CAHS D. In osmotically shocked cells expressing mVenus alone, we observed a decrease in cell volume of ∼50%, which was consistent with cells expressing our non-gelling mVenus:FL_Proline variant (Fig. 6C, Fig. S6G). By contrast, in osmotically shocked cells expressing gelling proteins mVenus:CAHS D and mVenus:2X Linker, we observed a reduction in volumetric change of only ∼1/3rd, which is significantly less than volume changes observed in mVenus and FL_Proline expressing cells (Fig. 6C).

These results suggest that CAHS D, and its variants, are able to better resist osmotically induced volumetric changes, but that this behavior requires the proteins to be in their condensation state.

To assess the degree to which cell viability correlates with the ability of a CAHS protein to reduce volumetric change during osmotic stress, we plotted these values and performed correlation analysis (Fig. 6D). While a mild correlation is observed (R^2^ = 0.55), this comparison indicates that FL_Proline (our non-gelling variant) protects cells more than expected, and 2X Linker (our gelling variant) protects cells less than expected, for the ability to reduce volumetric change to be the only protective property of these proteins (Fig. 6D).

### Gelling variants of CAHS D slow metabolism, which correlates with increased cellular survival, during osmotic shock

The observation that the reduction of volumetric change alone cannot fully explain the protective capacity of CAHS D and its variants during osmotic shock prompted us to examine additional potential mechanisms of protection. *In vitro*, CAHS D stabilizes enzymes and this is thought to be linked to the ability of CAHS D to slow molecular motions (Fig. 1F-H) [9,15,18]. We wondered if slowed molecular motions and stabilization displayed by CAHS D *in vitro* carries over to its *in vivo* function(s) and whether or not CAHS D might be inducing biostasis in cells.

To assess the ability of CAHS D to induce biostasis in cells and the necessity of gelation in this process, we monitored the metabolic activity of cells expressing mVenus, CAHS D, 2X Linker, and FL_Proline during osmotic shock using two different metabolic assays (Fig. S6H & I). During osmotic shock, we observed a statistically significant decrease in metabolism in cells expressing our two gelling proteins (CAHS D and 2X Linker) relative to cells expressing mVenus (Fig. 6E). By contrast, metabolism of cells expressing non-gelling FL_Proline was unchanged compared to the mVenus control (Fig. 6E). Correlation of metabolic rates with the degree of cellular survival during osmotic shock showed a robust relationship (R^2^ = 0.95) between the degree of reduced metabolism and the survival of stressed cells (Fig. 6F).

These results suggest that CAHS D may protect cells through the induction of biostatic slowdown of metabolism, which requires, and is induced by, condensate formation.

### Biostasis is reversible in CAHS D expressing cells

*In vitro*, CAHS D gels are highly soluble and go back into solution rapidly (Fig. 1C). We wondered if the same were true *in vivo* and, if so, whether the slowing of cell metabolism observed with condensed CAHS D or 2X Linker might also be reversible.

To assess this, we imaged cells recovering from osmotic stress (Fig. 6G). Consistent with *in vitro* results (Fig. 1C), mVenus:CAHS D in cells was observed to transition from the condensed state back into solution upon recovery from osmotic shock (Fig. 6G). Interestingly, this robust reversal out of the condensation state upon recovery from osmotic shock was much slower for mVenus:2X Linker expressing cells (Fig. 6G & H), consistent with a lack of resolubility of the 2X Linker variant *in vitro* (Fig. S5C).

In addition to observing the reversibility of the condensed state of mVenus:CAHS D in cells recovering from osmotic shock, we monitored the reversibility of metabolic depression. While metabolic activity of cells expressing mVenus:CAHS D and mVenus:2X Linker were significantly lower than cells expressing mVenus or mVenus:FL_Proline during osmotic shock (Fig. 6E), recovering cells expressing mVenus:CAHS D had metabolisms identical to those of recovering mVenus or mVenus:FL_Proline expressing cells (Fig. 6I, Fig. S6H-J). However, the metabolism of cells expressing mVenus:2X Linker remained depressed during recovery, mirroring the inability of 2X Linker to decondense (Fig. 6I, Fig. S6J, Fig. S5C). Consistent with the idea that condensation affects metabolism, cells expressing FL_Proline had metabolic rates identical to control cells (Fig. 6I, Fig S6J).

Combined, these data show that condensation and associated biostasis is reversible for wildtype CAHS D, but that this reversibility is compromised in our 2X Linker variant.

## Discussion

CAHS D is an intrinsically disordered protein from the tardigrade *H. exemplaris* that is necessary for this animal to successfully enter into and emerge from desiccation-induced biostasis [9]. Here we examine the sequence features and mechanisms underlying the ability of CAHS D to condense into a gel, and the consequences of condensation *in vivo* for mediating osmotic stress tolerance in cells. We find that the same sequence features drive *in vitro* gel formation also promote *in vivo* condensation of CAHS D. *In vivo* condensing and non-condensing forms of CAHS D promote cellular survival, however only the condensed forms help minimize volumetric change associated with osmotic shock. In addition, CAHS D condensation induces biostasis as evidenced by the slowing of cellular metabolism, which correlates with increased survival during osmotic stress. Finally, just as tardigrades are able to recover from their biostatic state, in cells, CAHS D decondenses upon return to normal osmotic conditions resulting in a return of metabolism to control levels. This study provides insight into how tardigrades, and potentially other desiccation-tolerant organisms, survive drying by making use of biomolecular condensation. Beyond stress tolerance, our findings provide an avenue for pursuing technologies centered around the induction of biostasis in cells and even whole organisms to slow aging and enhance storage and stability.

### Mechanism of CAHS D gel and condensate formation

Previous reports on the gelation of CAHS proteins have focused on mechanisms exclusively involving the association of helices found within the central Linker Regions of these proteins [19–21]. Here we combine molecular, biochemical, and biophysical experiments with computational analyses of CAHS D to reveal that while its Linker Region is required for gelation it is not sufficient. Rather, all three *N*-terminal, Linker, and *C*-terminal Regions, are required for gelation of this protein. Furthermore, cross-linking and predictive modeling suggests that antiparallel association of the linker results in dimers of CAHS D with heterotypic ends to promote further assembly. Indeed, CAHS D variants with homotypic termini do not form gels. Furthermore, the observation that the diameter of gelled fibers formed by wildtype CAHS D and a 2X Linker variant are statistically identical negates the idea that gels form through helical stacking.

Delving deeper into the mechanisms underlying gelation of CAHS D, we find that helicity and amphipathicity of the Linker Region is tuned to promote gelation. Also, β-β interactions between the terminal regions of CAHS D increase and are required for gelation. Based on these observations and coupled with predictions, we arrive at what we believe is the most parsimonious model of gelation given the current data. In this model, antiparallel association of Linker Regions mediates the association of two CAHS D molecules resulting in a staggered dimer with heterotypic ends. These dimers associate further into bundled structures with a diameter of ∼9.5 nm before associating further via end-to-end polymerization.

Importantly, measurements of fiber diameter from three different experiments (SAXS, AFM, and SEM) are in good agreement that CAHS D fibers have diameters ∼9 – 9.5 nm. While SEM imaging of CAHS D and 2X Linker show gel fibers with diameters of ∼12 nm, it should be remembered that SEM is known to minimally add ∼2 nm to surface thickness [23]. These measurements of fiber thickness inform our model of how dimers of CAHS D could come together to form such structures. However, equally important as the geometry of the surface of the fibers are the internal dimensions. SAXS analysis on CAHS D gels measures a slight change in the average size of internal void spaces within a fiber from 2.6 to 2 nm as the gel condenses. The presence of mesh structure within the fiber could be of importance, since filling a space with partially hollow fibers is more efficient than doing so with solid assemblies.

### Do CAHS gels represent a novel, stress-inducible cytoskeletal network?

By its most basic definition the cytoskeleton is a system of intracellular filaments or fibers that helps maintain a cell’s shape and/or organization [65]. Typically actin filaments, microtubules, and intermediate filaments are considered the three major classes of cytoskeletal components, but other cytoskeletal components exist, such as the filamenting temperature-sensitive mutant Z (ftsZ) protein, a prokaryotic functional homolog of tubulin involved in cytokinesis during cell division or spetins, which have been dubbed the ‘fourth component of the cytoskeleton’ [66,67]. Other common attributes of cytoskeletal networks are that they are polymers and often made from (homo)monomers (e.g., f-actin or β-tubulin) and that they can be dynamically assembled and disassembled [65].

Our proposed model of gelation shares similarities to that proposed for intermediate filaments (IFs; Fig. 7A). In IF assembly, two copies of an IF protein form parallel coiled-coil dimers, and two of these associate to form staggered antiparallel tetramers. Eight such tetramers then assemble into a bundle, which supercoils to form a unit-length filament (ULF) with a diameter of ∼8 –10 nm (Fig. 7A). These ULFs then further polymerize through end-to-end interactions between homotypic headgroups to form IFs [65,68].

**Figure 7.**
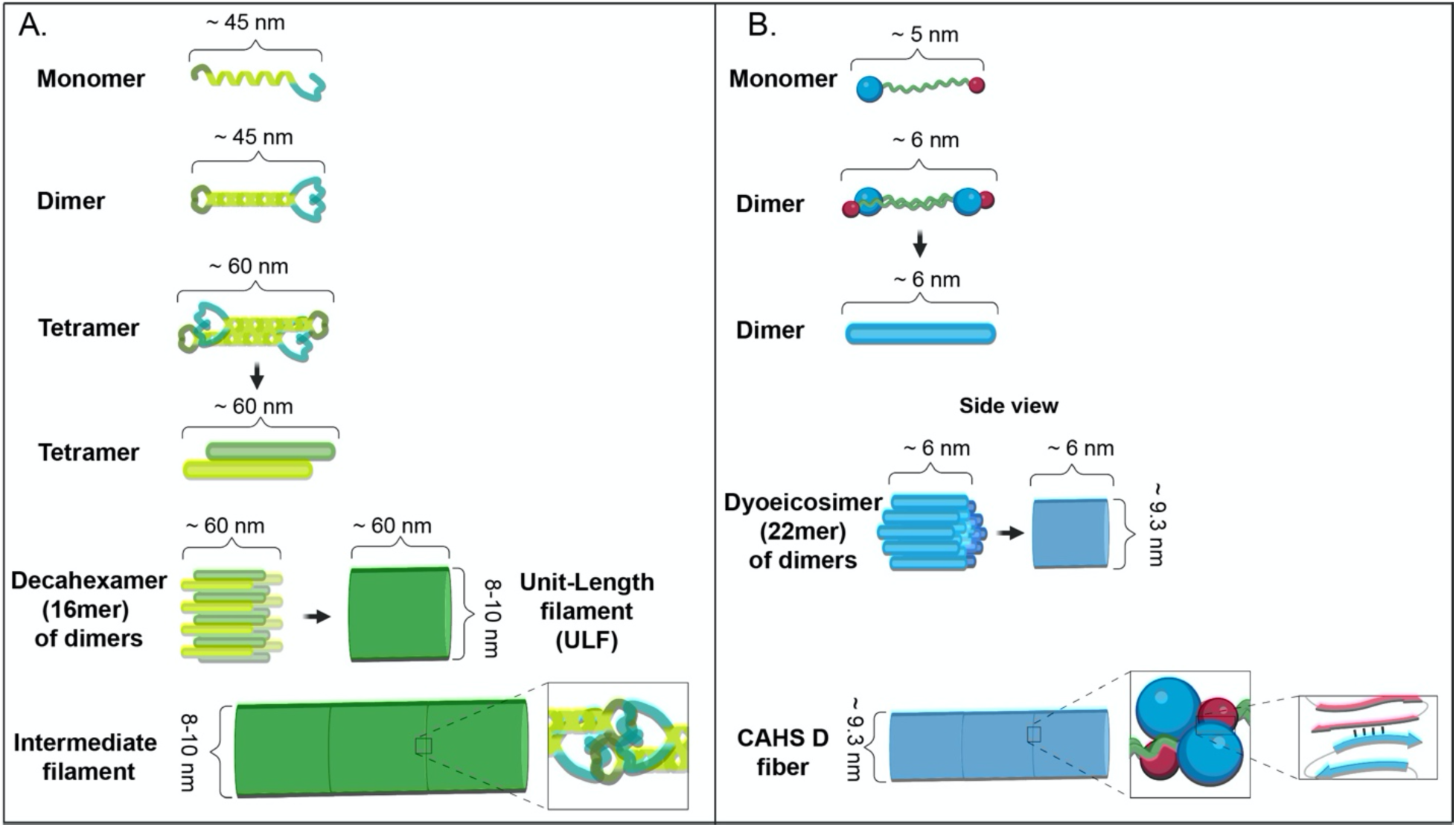
Comparisons of proposed mechanisms of intermediate filament and CAHS D fiber formation. **A)** Schematic representation of intermediate filament formation (adapted from [76]). **B)** Schematic representation of proposed mechanism of CAHS D fiber formation.

The similarities between CAHS D fibers and IFs extend beyond the mechanism of their formation. Indeed, there appear to be functional similarities between CAHS D fibers and IFs as well. IFs play diverse roles in cells, including the maintenance of cellular stiffness, organization, and integrity [65,68]. Here we show that CAHS condensates help maintain the volumetric integrity of cells during osmotic shock (Fig. 6C, Fig. S6G). In addition, a previous report demonstrates that CAHS proteins help mediate cellular stiffening [20]. Thus, due to shared similarities in assembly and function, we propose that CAHS proteins in part help protect cells through the induction of a novel, intermediate filament-like cytoskeletal network.

While IFs are typically thought of as less dynamic than actin filaments and microtubules there are examples of highly dynamic IF processes [69]. As their name implies, intermediate filaments were first distinguished from other cytoskeletal networks in that the fibers they form have an *intermediate* diameter (∼10 nm), between that of actin filaments (7 nm) and microtubules (25 nm) [70]. Given that CAHS proteins do not share sequence homology with *bona fide* IF proteins, but do share similarities in terms of assembly mechanism, dimensions, and function, we take the conservative view that CAHS D condensates represent a novel intermediate filament-like cytoskeletal network. Importantly, CAHS condensation is dynamic, inducible, and reversible, making them distinct from the current models for IF assembly.

### CAHS D as a biostasis-inducing protectant

While CAHS D condensation is associated with increased cellular survival, a non-condensing variant of CAHS D (FL_Proline) also promotes survival, albeit at significantly lower levels. This observation implies that polymerization into a novel cytoskeleton is not the only mechanism of protection by which CAHS D functions in cells during osmotic stress.

Induction of biostasis, the slowing down or halting of biological processes, has long been implicated in desiccation tolerance. This idea is encapsulated by both the molecular shielding [15,71,72] and vitrification [9,15,24,72] hypotheses. In the first, protectants act as molecular shields, getting in the way of aggregation prone proteins slowing or altogether preventing their interactions [15,18,71,73]. In the second, a super-viscous state is thought to be induced that ultimately forms an amorphous (vitrified) solid, preventing both large and small scale molecular motion such as diffusion of molecules and protein unfolding [9,15,24,72,74,75]. While vitrification is not sufficient to confer protection during desiccation (many non-protective molecules are known to vitrify) it is generally accepted as a requirement to survive in the dry state [24]. CAHS proteins, as well as tardigrades, have been shown to vitrify [9,56,75], which is not surprising given the capacity of CAHS proteins are disordered and form gels, which themselves are often amorphous solids [57,58].

Our observation that reduced cellular metabolism is a feature of osmotically shocked cells expressing variants of CAHS D that can condense (*in vivo*) or gel (*in vitro*) and that *in vivo* this correlates strongly with survival, suggests that induction of biostasis is a mechanism underlying CAHS D mediated protection and is specific to the protein’s capacity to form condensates. Notably, while osmotically stressed cells expressing our 2X Linker variant do show slowed metabolism, returning these cells to normal osmotic conditions does not lead to recovered metabolism. This continued depression of metabolic rates in these cells is mirrored by the 2X Linker protein’s diminished capacity to resolvate both *in vitro* and *in vivo* (Fig S5C, Fig 6G & H).

Thus, our study provides the first empirical evidence for not one, but two, condensation-specific roles for CAHS D: the maintenance of cell volume, and the induction of reversible biostasis during osmotic shock.

Our work advances our fundamental knowledge of how tardigrades survive extreme abiotic stress by providing a holistic and mechanistic model for the formation of a novel stress-inducible cytoskeleton composed of CAHS D, which promotes cellular survival through the maintenance of cell volume as well as through the induction of reversible biostasis. Beyond tardigrades, our work provides avenues for pursuing novel cell-stabilization/anti-aging technologies as well as provides insights into how CAHS proteins from tardigrades might be used and/or engineered to stabilize sensitive pharmaceuticals.

## Methods

### Cloning

All variants and wild type CAHS D except GS variants were cloned into the pET28b expression vector using Gibson assembly methods. Primers were designed using the NEBuilder tool (NEB), inserts were synthesized as gBlocks and purchased from Integrated DNA Technologies (IDT). Clones were propagated in DH5α cells available from NEB (Catalog C2987H).

GS variants were designed by replacing the Linker region of CAHS D with glycine-serine (GS) repeats of 25, 53 and 105 amino acids. The constructs were cloned after NotI in pMalC6T (NEB) by Twist Bioscience. Clones were propagated in DH5α cells available from NEB (Catalog C2987H).

No hydro, NAH, and RAH sequences were designed using sequence analysis tools built into sparrow (https://github.com/idptools/sparrow/) and GOOSE (https://github.com/idptools/goose).

### Protein Expression

Expression constructs were transformed in BL21 (DE3) *E*. *coli* (Catalog C2527H, NEB) and plated on LB agar plates containing 50 µg/mL Kanamycin for constructs in pET28b and containing 100 µg/mL Ampicillin for constructs in pMalC6T. At least 3 single colonies were chosen for each construct and tested for expression.

Large-scale expression for constructs in pET28b was performed in 1L LB/Kanamycin cultures, shaken at 37⁰C (Innova S44i, Eppendorf) until an OD 600 of 0.6, at which point expression was induced using 1 mM IPTG. Protein expression was continued for four hours, after which cells were collected at 4000 xg for 30 minutes at 4°C. Cell pellets were resuspended in 5 mL of resuspension buffer 20 mM tris, pH 7.5, 30 µL protease inhibitor (Catalog P2714-1BTL,Sigma Aldrich). Pellets were stored at −80⁰C.

Large-scale expression for constructs in pMalC6T was performed in 1L LB/Ampicillin supplemented with 0.2% glucose cultures, shaken at 37⁰C (Innova S44i, Eppendorf,) until an OD600 of 0.5, at which point expression was induced using 0.3 mM IPTG. Protein expression continued for four hours, after which cells were collected at 4000 xg for 30 minutes at 4°C. Cell pellets were resuspended in 10% w/v of resuspension buffer 20 mM tris, 200mM NaCl, 1mM EDTA, pH 7.4, 30 µL protease inhibitor (Catalog P2714-1BTL, Sigma Aldrich). Resuspended pellets were stored at −80⁰C until further use.

### Protein Purification

Purification for constructs in pET28b largely follows the methods in Piszkiewicz et al., 2019 [14]. Bacterial pellets were thawed and heat lysis was performed. Pellets were boiled for ten minutes and allowed to cool for 10 minutes. All insoluble components were removed via centrifugation at 13,000 xg at 10°C for 30 minutes. The supernatant was sterile filtered with 0.45 µm and 0.22 µm syringe filters (EZFlow Syringe Filter, Cat. 388-3416-OEM). The filtered lysate was diluted 1:2 in purification buffer UA (8 M Urea, 50 mM sodium acetate, pH 4). The protein was then purified using a cation exchange HiPrep SP/HP 16/10 (Cytiva,) on the AKTA Pure 25L (Cytiva), controlled using the UNICORN 7 Workstation pure-BP-exp (Cytiva). Variants were eluted using a gradient of 0-50% UB (8 M Urea, 50 mM sodium acetate, and 1 M NaCl, pH 4), over 20 column volumes.

Fractions were assessed by SDS-PAGE and pooled for dialysis in 3.5 kDa MWCO dialysis tubing (SpectraPor 3 Dialysis Membrane, catalog 132724, Sigma Aldrich). For all variants in pET28b except CLC, pooled fractions were dialyzed at 25°C for four hours against 2 M urea, 20 mM sodium phosphate at pH 7.0, then transferred to 20 mM sodium phosphate at pH 7 overnight. This was followed by six rounds of 4 hours each in Milli-Q water (18.2 MΩcm). Dialyzed samples were quantified fluorometrically (Qubit 4 Fluorometer, catalog Q33226, Invitrogen), aliquoted in the quantity needed for each assay, and lyophilized (FreeZone 6, Labconco) for 48 hours, then stored at −20°C until use. CLC was dialyzed in 2 M urea, 20 mM Tris at pH 7 for four hours, followed by 6 rounds of 4 hours each in 20 mM Tris pH 7. CLC samples were quantified using Qubit4 fluorometer as described, concentrated using spin-concentrators (Catalog UFC900324, Millipore-Sigma) to the desired concentration and used immediately.

Purification of the GS constructs in pMalC6T was performed following the NEB NEBEXpress MBP Fusion and Purification System protocols (Catalog E8201S, NEB). Pellets were thawed in cold water and lysed by sonication for 10 minutes, using 40% amplitude and 30 seconds on/30 seconds off cycles on ice. All insoluble components were removed via centrifugation at 13,000 xg at 10 °C for 30 minutes. The supernatant was filtered with 0.45 μm and 0.22 μm syringe filters (EZFlow Syringe Filter, Cat. 388-3416-OEM). The proteins were then purified by affinity chromatography using amylose resin (Catalog E8021L, NEB). The amylose resin was washed and pre-equilibrated in 20 mM tris, 200 mM NaCl, 1 mM EDTA, pH 7.4 buffer before the supernatant from the cell extract was loaded. After the supernatant was loaded the resin was washed with 12 column volumes of 20 mM tris, 200 mM NaCl, 1 mM EDTA, pH 7.4 buffer.

MBP and the GS linkers are fused by a polylinker containing a TEV protease recognition site for easy removal of the MBP-tag. Cleavage of GS linkers from MBP was done in column using 1 unit of TEV protease (Catalog P8112S, NEB) for every 2 µg of fusion protein in 1X TEV buffer (Catalog P8112S, NEB) for 16h at 30°C.

GS proteins were eluted as flow-through, using 2 column volumes of 20 mM tris, 200 mM NaCl, 1 mM EDTA, pH 7.4 buffer to wash off the proteins from the resin.

TEV Protease and MBP contain polyhistidine tags at their N-termini and they can be removed from the cleavage reaction by immobilized metal affinity chromatography, such as Nickel or Cobalt resin, TEV protease and any free MBP or uncleaved MBP fusion were further removed using a cobalt resin (Catalog 786-402, G-Biosciences).

Purified GS proteins were assessed by SDS-PAGE and pooled for dialysis in 3.5 kDa MWCO dialysis tubing (SpectraPor 3 Dialysis Membrane, catalog 132724, Sigma Aldrich). Proteins were dialyzed at 25 °C for 4 hours against 25 mM Tris-HCl, 150 mM NaCl, pH 7.4, then transferred to 25 mM Tris-HCl, 50 mM NaCl, pH 7.4 overnight. This was followed by 4 rounds of 4 hours each in Milli-Q water (18.2 MΩcm). Dialyzed samples were quantified fluorometrically (Qubit 4 Fluorometer, catalog Q33226, Invitrogen), aliquoted in the quantity needed for each assay and lyophilized (Catalog 7752021, Labconco FreeZone 6) for 48 hours, then stored at −20 °C until use.

### Visual assay for concentration and temperature dependence of gelation

Quantitated and lyophilized protein samples were transferred as powder into NMR tubes (Catalog WG-4001-7, Bel-Art Products) and resuspended in 500 µL of water to final concentrations of 5 g/L, 10 g/L, 15 g/L, and 20 g/L. Samples were left at room temp for 5 minutes to solubilize. If solubilization was not occurring (as determined visually), samples were moved to 55°C for intervals of 5 minutes until solubilized. If solubilization was not progressing at 55°C after 10 minutes of heating (as determined visually), then samples were transferred 95°C for 5-minute intervals until fully solubilized.

Solubilized proteins were transferred from heat blocks to the bench and left at ambient temperature for 1 hour. Tubes were then loaded horizontally into a clamp holder and photographed. Gelation was visually assessed by the degree of solidification or flow of the sample in the NMR tube.

### Visual assay for gel resolubility assessment

After 1 hour at ambient temperature, proteins that had been found to form gels were transferred to a 55°C heat block for 1 hour to thermally resolubilize them. After an hour, samples were returned to the photographic clamp holder and imaged immediately to confirm that gelation had been disrupted. Samples were placed upright on the bench at ambient temperature for 1 hour to gels reform upon cooling to room temperature.

To assess resolubility, samples of 10 g/L CAHS D and 2X LR were prepared as described above. Buffer (Tris, 20 mM pH 7) was added to the gels to bring the final concentration of solvated CAHS D and 2X LR to 2.5 g/L, which is below the gelation point of both proteins. Samples were photographed before and immediately after addition of buffer, vortexed for 5 seconds, and left to resolubilize. For heat resolubilization resolvated samples were moved to a 55°C heat block and the time for solubilization was recorded.

### Scanning electron microscopy, critical point drying, and image analysis

Protein samples were heated to 95°C for 5 minutes and 50 µL of each sample were transferred to microscope coverslips. Samples were fixed in a 2.5% glutaraldehyde / 4% paraformaldehyde solution for 10 minutes. Samples were then dehydrated in an ethanol series going from 25%, 50%, 75%, 3x 100% with each incubation time being 10 minutes. Dehydrated samples were prepared for imaging using critical point drying (Tousimis Semidri PVT-3, SGC Equipment) and supporter coating (108 Auto, Cressington Scientific Instrument). Imaging was performed on a Hitachi S-47000 scanning electron microscope. Image analysis for fiber diameter and ULF lengths was carried out using FIJI (V. 2.3.0/1.52q).

### Lactate Dehydrogenase Protection Assay

LDH desiccation protection assay was performed in triplicate as described previously [9,14]. Briefly, CAHS D was resuspended in a concentration range from 20 g/L to 0.1 g/L in 100 µL resuspension buffer (25 mM Tris, pH 7.0). Rabbit Muscle L-LDH (catalog 10127230001, Sigma-Aldrich) was added to this at 0.1 g/L. Half of each sample was stored at 4°C, and the other half was desiccated for 17 hours without heating in a speedvac (OFP400, Thermo Fisher Scientific). Following desiccation all samples were brought to a volume of 250 µL with water. The LDH/CAHS D mixture was added 1:10 to the assay buffer (100 mM Sodium Phosphate, 2 mM Sodium Pyruvate, 1 mM NADH, pH 6). Enzyme kinetics were measured by NAD^+^ absorbance at 340 nm, on the NanodropOne (Thermo Fisher Scientific). The protective capacity was calculated as a ratio of NAD^+^ absorbance in desiccated samples normalized to non-desiccated controls. Sigmoidal curve has been added as a visual guide.

### FTIR Sample Preparation and Measurement

All samples were prepared by dissolving the CAHS D protein in D_2_O to avoid the overlapping of the amide I band of CAHS D (centered at 1650 cm^−1^) with the H_2_O bending mode (around 1640 cm^−1^) [77]. Gel samples were obtained by dissolving the lyophilized CAHS D protein at a concentration of 16 g/L in D_2_O heated at about 50 °C and gently stirred for 5 minutes. A volume of 10 µL of the CAHS D solution was deposited between two CaF_2_ windows, mounted into a Jasco MagCELL sample holder. Glassy samples were obtained by depositing a 38 µL drop of the heated CAHS D solution (16 g/L) in D_2_O on a 50 mm diameter CaF_2_ window. The sample was dried under N_2_ flow for 5 minutes and subsequently the optical window on which the glass had formed was inserted into a sample holder equipped with a second CaF_2_ window to form a gas-tight cylindrical cavity [78]. Different hydration levels of the CAHS D glass were obtained by an isopiestic method, i.e. by equilibrating the sample with saturated salt solutions providing defined values of relative humidity (RH), contained at the bottom of the sample holder cavity. The following saturated solutions in D_2_O were employed to obtain the desired RH [79] at 297 K: KNO_3_ (RH=95%), NaCl (RH=75%), Mg(NO_3_)_2_ (RH=53%), K_2_CO_3_ (RH=43%), MgCl_2_ (RH=33%), CH_3_COOK (RH=23%), and LiCl (RH=11%).

FTIR absorption measurements were performed at room temperature with a Jasco Fourier transform 6100 spectrometer equipped with a DLATGS detector. The spectra, in the range (7,000 - 1,000 cm^−1^) with a resolution of 2 cm^−1^, were acquired by averaging 10^3^ interferograms, using a standard high-intensity ceramic source and a Ge/KBr beam splitter.

### Amide hydrogen/deuterium exchange kinetics in a CAHS D dried matrix at different hydration levels

Amide H/D exchange can be followed by FTIR spectroscopy of the amide II band of proteins centered around 1550 cm^−1^ in H_2_O. Following H/D exchange the wavenumber of this mode is downshifted at 1450 cm^−1^ [80] (the so-called amide II’ band) Thus, the two bands are well separated in the spectrum.

Two samples of CAHS D protein in H_2_O (16 g/L) were extensively dried by equilibration within gas-tight holders at relative humidity RH=6% in the presence of NaOH·H_2_O, by using the isopiestic method described in the FTIR Sample Preparation and Measurement section. After equilibration, H/D exchange at RH=11% was started in one of the samples by placing saturated LiCl in D_2_O in the sample holder to create an 11% D_2_O atmosphere. H/D exchange at RH=75% was begun in the second sample by placing a saturated solution of NaCl in D_2_O in the holder. We have previously shown [81,82] that such an isopiestic approach for isotopic exchange is quite effective and rapid (hour time scale). In order to obtain the kinetics of H/D exchange (Fig. 1G, Fig. S1E), a series of FTIR spectra was then recorded in sequence, at selected time intervals following the start of H/D exchange. For each spectrum, only 100 interferograms were averaged to allow a sufficiently rapid acquisition (3 minutes). We evaluated the extent of amide H/D exchange from the area of the amide II’ band at 1450 cm^−1^ after subtraction of a straight baseline drawn between the minima on either side of the band [83].

### Measurement of conformational dynamics of photosynthetic reaction centers embedded in CAHS D glasses

The photosynthetic reaction center (RC) from the purple bacterium *Rhodobacter* (*Rb.*) *sphaeroides represents* an ideal model system to probe matrix effects on conformational protein dynamics. This membrane-spanning pigment-protein complex catalyzes the primary photochemistry of bacterial photosynthesis. Following photoexcitation, the primary electron donor (P) of the RC (a bacteriochlorophyll dimer situated near the periplasmic side of the protein) delivers an electron in about 200 ps to the primary quinone acceptor, Q_A_ (located 25 Å away from P and closer to the RC cytoplasmic side) thus generating the primary charge separated state, P^+^Q_A_^−^ (Fig. S1F). In the presence of o-phenanthroline, (an inhibitor which blocks electron transfer from Q_A_^−^ to Q_B_), the electron on Q_A_^−^ recombines with the hole on P^+^ by direct electron tunneling [26].

The kinetics of P^+^Q_A_^−^ recombination after a short (ns) flash of light provides an endogenous probe of the RC conformational dynamics. In fact, when the RC relaxation from the dark-adapted to the light-adapted conformation is hampered (e.g. at cryogenic temperatures [29], or by incorporation of the RC into dehydrated trehalose glasses [27]) the recombination kinetics are accelerated and become strongly distributed, mirroring the immobilization of the protein over a large ensemble of conformational substrates. Accordingly, charge recombination kinetics of confined RCs are described by a continuous distribution p(*k*) of rate constants *k* [27] (Fig. 1H, Fig. S1G & H):

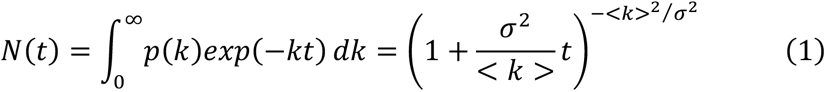

where N(t) is the survival probability of the P^+^Q_A_^−^ state at time t after photoexcitation, <*k*> is the average rate constant, and σ is the width of the rate distribution p. An increase in either or both parameters (<*k*>, σ) reflects inhibition of the RC internal dynamics, i.e slowing of relaxation from the dark- to the light-adapted conformation (<*k*>), or of the interconversion between conformational substates (σ) [27,28].

CAHS D : RC glassy samples were prepared by mixing *a* 78 µL volume of 76 µM RC purified from *Rb. sphaeroides* R26 in assay buffer (10 mM Tris, 0.025% LDAO, pH 8.0) with 64 µL of 16 g/L CAHS D protein in water, and 8 µL of 200 mM o-phenanthroline in ethanol. This mixture was deposited at the center of a 50 mm diameter CaF_2_ optical window and solid samples were obtained and equilibrated at RH by using the isopiestic approach described in the *FTIR Sample Preparation and Measurement* section.

The kinetics of P^+^Q_A_^−^ recombination after a laser pulse (Fig. 1H) were measured by time-resolved optical absorption spectroscopy, by recording the absorbance change at 422 nm, essentially as described in [84]. At each RH tested the residual water content of the CAHS D : RC matrices was determined from the area of the (*v*_2_+*v*_3_) combination band of water at 5155 cm^−1^ [28,78]. The water content of the matrices has been expressed as the mass ratio of water : dry matrix. To determine the mass of the dry matrix, we included the CAHS D protein, the RC, and the detergent belt of the RC formed by 289 LDAO molecules [78] plus 14 molecules of free LDAO per RC. The hydration isotherms obtained for dried CAHS D : RC and trehalose : RC matrices are essentially coincident and well fit the Hailwood and Horrobin equation [85] (Fig. S1I).

### Proteome-wide bioinformatics

The tardigrade proteome (tg.default.maker.proteins.fasta), taken from https://github.com/sujaikumar/tardigrade was used. The proteome file was pre-processed using protfasta (https://protfasta.readthedocs.io/), IDRs predicted with metapredict [86] and IDR kappa values calculated using localCIDER [34,35]. Metapredict identified 35,511 discrete IDRs distributed across 39,532 proteins in the tardigrade proteome.

### All-atom simulations

All-atom simulations were run using the ABSINTH implicit solvent model and CAMPARI Monte Carlo simulation engine (https://campari.sourceforge.net/). ABSINTH simulations were performed with the ion parameters derived by Mao et al., 2012 [87]. The combination of ABSINTH and CAMPARI has been used to generate experimentally-validated ensembles for a wide range of IDRs [88–90].

Simulations were performed for the full-length CAHS D protein starting from a randomly generated non-overlapping starting state. Monte Carlo simulations update the conformational state of the protein using moves that perturb backbone dihedrals, and sidechain dihedrals, and rigid-body coordinates of all components (including explicit ions).

Ten independent simulations were run for 150,000,000 steps each in a simulation droplet with a radius of 284 Å at 310 K. The combined ensemble generated consists of 69,500 conformations, with ensemble average properties computed across the entire ensemble where reported. Finally, we subsampled to analyze one in every 10 frames, providing us with an ensemble of 6950 frames total. All simulation analyses presented here were performed on this smaller ensemble of 6950 conformers, although equivalent results are obtained if the full 69,500 conformer ensemble is used. We reduced the ensemble size down to 6950 to make Bayesian reweighting via SAXS curves tractable, as to re-weight, each conformer in the ensemble must have a scattering curve calculated (discussed below).

Given the size of the protein, reliability with respect to residue-specific structural propensities is likely to be limited, such that general trends should be taken as a qualitative assessment as opposed to a quantitative description. We share this caveat especially in the context of future re-analysis of our simulations. Simulations were analyzed using SOURSOP and MDTraj [91,92]. Simulated scattering profiles were generated using FOXS [93].

### Polymer models

In addition to performing all-atom simulations, in Fig. 2 we reported distributions for the radius of gyration for two reference states: the Analytical Flory Random Coil (AFRC) and the self-avoiding random coil (SARC), also previously referred to as the Excluded Volume (EV) simulation in our previous work.

The AFRC model generates a sequence-specific Gaussian chain parameterized on all-atom simulations to generate an ensemble that matches the expected dimensions if CAHS D behaved like a Gaussian chain [33]. The SARC model is a numerically-generated ensemble obtained by performing all-atom simulations in which all interactions other than the repulsive component of the Lennard-Jones potential are turned off, resulting in a sequence-specific model of the excluded volume accessible to a polypeptide. For more details on the SARC model, see Holehouse et al. 2015, where we refer to it as the EV ensemble [94].

### Bayesian reweighting of simulations

The subsampled simulation ensemble (6,950 frames) was reweighted by the Bayesian/Maximum-Entropy (BME) method using a Python script from Bottaro and colleagues (https://github.com/KULL-Centre/BME) [32]. Per-frame scattering curves were calculated from simulation data using FOXS [93]. The goodness of fit between the calculated and experimental SAXS curves prior to reweighting is described by an initial reduced chi-squared (χ^2^) value of 11.25. The parameter θ is a scaling factor such that a lower θ value should yield a lower reweighted χ^2^ (better agreement with experimental data). However, as θ and χ^2^ decrease, so does the fraction of effective simulation frames (Φ) used in the reweighted ensemble. In this case, χ^2^ was largely unchanged at θ values of < 100, so we set θ = 50 to maximize the fraction of simulation frames reflected in the reweighted ensemble.

The reweighted ensemble shows a much better agreement with the SAXS data (χ^2^ = 1.71) and preserves ∼46% of the per-frame simulation data (Φ = 0.46). Note that no simulation frames are discarded in the reweighted ensemble, but the BME protocol gives each frame a new weight such that the sum of the weights of all frames equals 1. For ensuing simulation analysis using the reweighted ensemble, the per-frame weights were applied when calculating ensemble-averaged features including the radius of gyration, intramolecular distances, and secondary structure.

### Small-angle X-ray scattering

All SAXS measurements were performed at the BioCAT (beamline 18ID at the Advanced Photon Source, Chicago, IL). SAXS measurements on monomeric CAHS D were collected with in-line size exclusion chromatography (SEC-SAXS) coupled to the X-ray sample chamber to ensure the protein was monomeric. Concentrated protein samples were injected into a Superdex 200 increase column (Cytiva) pre-equilibrated in a buffer containing 20 mM Tris pH 7, 2 mM DTT, and 50 mM NaCl. Scattering intensity was recorded using a Pilatus3 1 M (Dectris) detector placed 3.5 m from the sample, providing a q-range from 0.004-0.4 Å-1. One-second exposures were acquired every two seconds during the elution. Data were reduced at the beamline using the BioXTAS RAW 1.4.0 software [95]. The contribution of the buffer to the X-ray scattering curve was determined by averaging frames from the SEC eluent, which contained baseline levels of integrated X-ray scattering, UV absorbance, and conductance. Baseline frames were collected immediately pre- and post-elution and averaged. Buffer subtraction, subsequent Guinier fits, and Kratky transformations were done using custom MATLAB (Mathworks) scripts or with custom Python code using the BioXTAS RAW 2.1.4 API (https://bioxtas-raw.readthedocs.io/en/latest/)

CAHS D samples were prepared for SAXS measurements by dissolving 5 mg/mL lyophilized CAHS D protein into a buffer containing 20 mM Tris pH 7 and 50 mM NaCl. Samples were incubated at 60°C for 20 minutes to ensure the sample was completely dissolved. Samples were syringe filtered to remove any remaining undissolved protein before injecting 1 mL onto the Superdex 200 column (Cytiva).

SAXS data for CAHS D gels were obtained by manually centering capillaries containing premade gels in the X-ray beam. Data was recorded as a series of thirty 0.2 second exposures, but only the first exposure was analyzed to minimize artifacts from X-ray damage. The final analyzed data was corrected for scattering from the empty capillary and a buffer containing capillary. CAHS D gel-containing samples were made by dissolving 100 mg/mL lyophilized protein in a buffer containing 20 mM Tris pH 7 and 50 mM NaCl. The sample was incubated for 20 minutes at 60°C to ensure the protein was completely dissolved. Double open-ended quartz capillaries with an internal diameter of 1.5 mm (Charles Supper) were used to make the samples. Dissolved protein was directly drawn into the capillary via a syringe. Concentration gradients were generated by layering the protein with buffer. Both ends of the capillary were then sealed with epoxy. Samples were allowed to cool for 5 hours prior to measurement. Because of the buffer layering method used to create the concentration gradient in gelled CAHS D samples, the precise concentration of the gel at any given point along the length of the capillary tube is not explicitly known. A concentration gradient can be defined from the bottom to the top of the capillary tube up as high to low CAHS D concentration. Data were collected along the concentration gradient by collecting data in 2 mm increments vertically along the capillary.

All data analysis was done using custom MATLAB (Mathworks) scripts. First, an effective concentration was calculated by assuming the maximum concentration was 100 mg/mL and scaling the remaining samples by the integrated intensity of the form factor. It should be noted that the actual concentration could be significantly less than 100 mg/mL in the maximum concentration sample. Data was fit to an equation containing three elements to describe the features apparent in the scattering data. The high-angle form factor was modeled using a Lorentzian-like function to extract the correlation length and an effective fractal dimension.

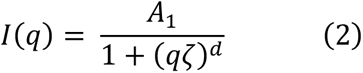

The correlation length is given by ζ and is related to the mesh size inside the fiber bundles seen in SEM images. The fractal dimension, *d*, is related to the density of the mesh. No clear correlation length was observed in the smallest angle data, and thus a power law was used to account for this component. The exponent *d*, is related to the nature of the interface inside and outside of the bundles.

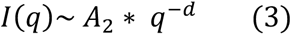

Finally, a Lorentzian peak was used to fit the diffraction peak that is apparent at higher concentrations. The width of the peak, B, appeared constant and was thus fixed so that the amplitude could be accurately extracted.

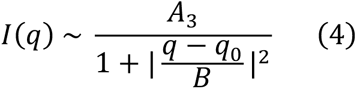

In all fit components, Ax is a scale factor.

### Ensemble Optimization Method

Ensemble Optimization Method (EOM) data was generated using ATSAS online’s EOM web portal (https://www.embl-hamburg.de/biosaxs/atsas-online/eom.php) [96,97]. Inputs for the simulation included a buffer subtracted SEC SAXS curve of CAHS D and the amino acid sequence of CAHS D. The terminal regions (residues 1-90 and 196-227) of CAHS D were described as broadly disordered and the linker region (residues 91-195) was described as broadly helical. No distance constraints were used. The average radius of gyration was determined from the EOM ensemble histogram using the following equation:

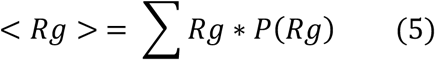

In which R_g_ is the value and P(R_g_) is the EOM-derived probability for that R_g_. The reported error is the standard deviation from the above average, calculated according to:

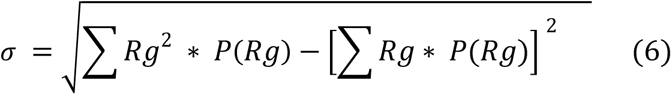

### CD Spectroscopy at non-gelling concentrations

Lyophilized protein constructs were weighed and dissolved in a 20 mM Tris-HCl (Fisher Bioreagents) buffer at pH 7.0. CD spectra were measured using a JASCO J-1500 CD spectrometer with 0.1 cm quartz cell (Starna Cells, Inc,) using a 0.1 nm step size, a bandwidth of 1 nm, and a scan speed of 200 nm/min. Each spectrum was measured 7 times and averaged to increase the signal-to-noise ratio. The buffer control spectrum was subtracted from each protein spectrum (Fig. S2K & L, Fig. S3A-D). The resulting spectra were deconvoluted to predict secondary structure contents using the lsq_linear function from the SciPy library. To do this, base spectra for α-helix, β-sheet, and random coil spectra (taken from [98]) were linearly fit to match the experimental data set.

### CD spectroscopy at gelling concentrations

To prepare stock solutions, freeze-dried proteins were dissolved in 20 mM MES buffer pH 6.0 and then pH was adjusted to 6.00±0.02 at 25.0±0.1 °C using pH Sensor InLab® Ultra-Micro-ISM (Mettler Toledo) calibrated before measurement. Gelled proteins were melted at 50 °C prior to pH measurement and adjustment, and then equilibrated at 25.0±0.1 °C.

CD data were collected on a JASCO J-810 spectropolarimeter fitted with a Peltier temperature controller (Jasco UK). A quartz cuvette with a light path of 0.1 cm was used. The samples were equilibrated in the CD instrument thermostated at 5 °C for 10 minutes prior to recording full spectra between 190 or 200 and 260 nm with a 1 nm step size, 100 nm·min^−1^ scanning speed, 1 nm bandwidth and 1 second response time. Then variable temperature experiments were performed by heating and cooling samples 5–90–5 °C at a rate of 30 °C/h whilst monitoring CD at 222 nm at 0.5 °C intervals followed by full spectra at 5 °C.

Baselines recorded using the same buffer, cuvette and parameters were subtracted from each dataset. The spectra were converted from ellipticities (deg) to mean residue ellipticities (MRE, (deg.cm^2^.dmol-1.res^−1^)) by normalizing for concentration of peptide bonds and the cell path length using the equation 7:

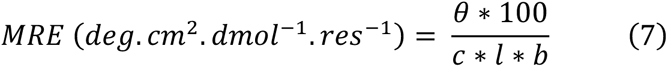

Where the variable θ is the measured difference in absorbed circularly polarized light in millidegrees, c is the millimolar concentration of the specimen, l is the path-length of the cuvette in cm and b is the number of amide bonds in the polypeptide, for which the N-terminal acetyl bond was included but not the C-terminal amide.

Fraction helix (%) was calculated from MRE at 222 nm using the following equation 8

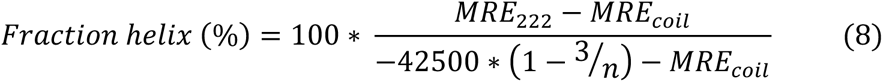

Where *MRE_coil_* = 640-45T; T is the temperature in °C; and n is the number of amide bonds in the sample (including the C-terminal amide) [99].

### Helical Wheel Plot

Helical wheel plots were generated using the heliQuest sequence analysis module. CAHS D linker sequence 126-144 was used, with the A-helix option chosen as the helix type.

### Differential Scanning Calorimetry (DSC) method

Prior to DSC measurement, samples were prepared in Eppendorf tubes at the desired concentration by adding protein into water. Protein solutions were then vortexed and heated to 55°C for 5 minutes to ensure proper mixing and that samples were completely solubilized. 40 µL of solubilized sample was hermetically sealed into a previously massed pair of DSC aluminum hermetic pan and hermetic lid (Catalog 900793.901 and 901684.901, respectively, TA instruments). The sample mass was determined after the sample was sealed within the pan and lid. The sealed samples were then run on a TA DSC2500 instrument. Trios software (TRIOS version #5.0.0.44608, TA Instruments) provided by TA Instruments was used to perform analysis of the DSC data. Details for DSC methods for specific figures are listed below:

> *Fig 1A* DSC method

> The heating run for different concentrations of CAHS D samples consisted of samples being equilibrated at 20 °C and then heated to 80 °C at a 5 °C per minute ramp.

> The heating and cooling run consisted of the samples starting at the stand-by temperature of 20 °C, then heated to 80 °C at a 5 °C per minute ramp, then cooled to 20 °C at a 5 °C per minute ramp, then held for a 10-minute isothermal hold at 20 °C, and finally the samples were heated to 80 °C at a 5 °C per minute ramp.

> *Fig 3B* DSC method

> The heating runs consisted of the samples being equilibrated at 10 °C and then heated to 95 °C at a 5 °C per minute ramp.

> *Fig 4B* DSC method

> The heating runs consisted of the samples being equilibrated at 20 °C and then heated to 80 °C at a 5 °C per minute ramp.

### Sequence analysis

In Fig. 2, sequence analysis was performed using sparrow (https://github.com/idptools/sparrow) and disorder prediction was performed using metapredict [100]. Sequence to radius of gyration predictions from ALBATROSS were performed using the metapredict webserver (https://metapredict.net/), which uses sparrow (https://github.com/idptools/sparrow) [31].

### Coiled-Coil predictions

Waggawagga web-based user interface was used to predict coiled-coil propensity on CAHS D and variants. Waggawagga is designed to provide a direct schematic comparison of many coiled-coil prediction tools. This interface allows the comparative analysis of six coiled-coil classic prediction programs (Marcoil, Multicoil, Multicoil2, Ncoils, Paircoil, Paircoil2) and three oligomerization state prediction programs (Scorer, PrOCoil and LOGICOIL). In addition, Multicoil2 distinguishes dimers, trimers and non-coiled-coil oligomerization states. These tools can be run in any combination against single or multiple query sequences [101].

For our comparative analysis we run Marcoil, Multicoil2, Ncoils and Paircoil2 classic prediction programs on a 14 aa window length with Scorer and LOGICOIL oligomerization state prediction programs.

### Crosslinking assays

BS3 (bis[sulfosuccinimidyl] suberate) (Catalog A39266, Thermo Scientific) a 11.4 Å arm amine ester crosslinker was used following manufacturer’s instructions in the crosslinking assays.

In figures 3F-I, a solution of 1 mg/mL of each protein was crosslinked with increasing amounts of BS3 (0.8 mM to 8 mM) for 30’ at room temperature. A sample with no crosslinker was used as control for the monomeric state of the proteins.

In figure S5D, we looked at the oligomeric state after buffer resolvation and after heat resolvation of CAHS D gels and crystalline solids. To obtain a CAHS D gel, a 20 mg/mL gel sample was prepared as described in the temperature and concentration dependent resolubility assay section. To prepare the crystalline solids, a gel formed with 20 mg/ml CAHS D was further dessicated in a speedvac (OFP400, Thermo Fisher Scientific) for 16 hours. The crosslinking assay was performed as follows; 2 mg/mL of CAHS D solution was crosslinked with 1 mM BS3 (lane 3) for 30’ at room temperature. As control for monomeric state, a sample of 2 mg/mL CAHS D without crosslinker was used (lane 1). In lane 3 a CAHS D gel formed at 20 mg/mL protein was resolubilized adding buffer to 2 mg/mL concentration and crosslinked with 1 mM BS3 for 30’ at room temperature. In lane 4 the buffer resolubilized gel was further heat resolubilized at 55°C until was clear and then crosslinked with 1 mM BS3 for 30’ at room temperature. In lanes 5 and 6 the same experiment was performed as in lanes 3 and 4 but using CAHS D crystalline solids.

After crosslinking assays, 15 µL of each sample was run in denaturing SDS-Page gels and stained with Coomassie blue to visualize CAHS D and the variants oligomeric states.

### 2D-IR

A detailed description of the setup used to measure the 2D-IR spectra can be found in [102]. Briefly, pulses of a wavelength of 800 nm and with a 40 fs duration were generated by a Ti:sapphire oscillator and further amplified by a Ti:sapphire regenerative amplifier to obtain 800 nm pulses at a 1 kHz repetition rate. These pulses were then converted in an optical parametric amplifier to obtain mid-IR pulses (C20 μJ, C6100 nm) that had a spectral full width at half-maximum (FWHM) of 150 cm^−1^. The beam was then split into probe and reference beams (each 5%) and a pump beam (90%) that was aligned by a Fabry–Pérot interferometer. The pump and probe beams overlapped in the sample in an C250 μm focus. The transmitted spectra of the probe (*T*) and reference (*T*_0_) beams with the pump on and off were then recorded after dispersion by an Oriel MS260i spectrograph (Newport, Irvine, CA) onto a 32-pixel mercury cadmium telluride (MCT) array. The probe spectrum was normalized to the reference spectrum to compensate for pulse-to-pulse energy fluctuations. The 2D-IR signal was obtained by subtracting the probe absorption in the presence and absence of the pump pulse. Parallel and perpendicular 2D-IR spectra were recorded by rotating the pump beam at a 45° angle with respect to the probe beam and selecting the probe beam component that was either perpendicular or parallel to the pump beam using a polarizer after the sample. To minimize pump-scattering contributions, we measured the average of two photoelastic modulator (PEM)-induced pump delays, such that the interference between the scattered pump beam and the probe beam had a 180° phase in one delay with respect to the other delay.

### FTIR analysis of the amide I’ band in CAHS D gels and glasses at increasing hydration

With the aim of assessing CAHS D secondary structure and its possible changes upon vitrification and dehydration we performed a FTIR spectral analysis of CAHS D amide I band in D_2_O (amide I’ band) [80]. Any particular secondary structure absorbs predominantly in a specific range of the amide I’ region; due to overlapping, however, the resulting amide I’ band is scarcely structured, and band narrowing procedures are necessary to resolve the various secondary structure components [103]. The number and positions of overlapping spectral components have been evaluated from the minima and maxima of the second and fourth derivatives, respectively, of the amide I’ band [104]. Second and fourth derivative spectra were calculated using the i-signal program (version 2.72) included in the SPECTRUM suite (http://terpconnect.umd.edu/~toh/spectrum/iSignal.html) written in MATLAB language. A Savitsky-Golay algorithm was employed to smooth the signals and calculate the derivatives. For all the amide I’ bands analyzed, in gel and glassy samples, both the second and fourth derivative spectra were consistently dominated by three minima and maxima, respectively, suggesting the presence of three spectral components, with peak wavenumbers in the intervals 1619-1628 cm^−1^ (ascribed to β-β contacts), 1644-1648 cm^−1^ (unordered/helical regions), and 1676-1688 cm^−1^ (β-β contacts) [80,103]. Amide I’ bands were decomposed into the identified three Gaussian components by using a locally developed least-squares minimization routine [28] based on a modified grid search algorithm [105]. Confidence intervals for the best-fit parameters were evaluated numerically, as previously detailed [106]. The fractional area of each component band was taken as the relative content of the corresponding secondary structures (Fig. 4E).

### Thioflavin T Assay

Proteins were dissolved in phosphate buffered saline, pH 7.2 (Sigma-Aldrich) Thioflavin T was prepared in Dimethyl Sulfoxide (Sigma-Aldrich) and diluted to 20 µM in PBS or Tris for use in the assay. Thioflavin T was prepared fresh for each assay at a concentration of 400 µM in DMSO, then diluted to 20 µM in PBS/Tris. Twenty five microliters of protein and ThT in PBS/Tris were combined in a 96-well plate (Catalog 3596, Costar, Fisher Scientific) and incubated for 15 minutes at room temperature in the dark. Fluorescence was measured using a plate reader (Spark 10M, Tecan) with an excitation at 440 nm, emission was collected at 486 nm. Assays performed using Tris rather than PBS were those measuring CLC due to solubility issues, which was resuspended in 20 mM Tris buffer pH 7.5. For these assays, Thioflavin T was also diluted in this buffer. All proteins and controls were assayed in triplicate.

### Cell line generation and maintenance

Full-length CAHS D (Uniprot P0CU50, CAHS 94063) and CAHS D variants FL_proline and 2X Linker with mVenus proteins fused in their N-termini were cloned into pTwist-cmv-WPRE-Neo between HindIII and BamHI by Twist Bioscience. 1 µg of plasmid DNA expressing mVenus:CAHS D, mVenus:FL_ Proline and mVenus:2x linker proteins were transfected into HEK293 (Catalog CRL-1573, ATCC) cells with lipofectamine 3000 transfection reagent (Catalog L3000008, Thermo Fisher Scientific). 24h post-transfection, cells that had successfully integrated mVenus, mVenus:CAHS D, mVenus:FL_Proline and mVenus:2X Linker were selected with 0.7 µg/µL G418 (Catalog G64500, Research products international). Cells were passed 2x before expanding and flash-cooling. Stable cell lines were maintained by supplementing Dulbecco’s modified Eagle’s medium (DMEM) (Catalog 10567014, Gibco) with 10% fetal bovine serum (FBS) (Catalog 900-108, GeminiBio Products), 1% penicillin/streptomycin (Catalog 400-109, Gemini Bio products) and 0.3 µg/µL G418 (Catalog G64500, Research products international) at 37 °C with 5% CO2 atmosphere.

### Fluorescence imaging

8-well glass bottom dishes (Catalog 80826, Ibidi) were pre-coated with 0.1 mg/mL poly-D lysine (Catalog 3890401, Thermo Fisher) for 1 h at 37°C. Cells expressing mVenus, mVenus:CAHS D, mVenus:FL_Proline and mVenus:2X Linker were seeded at a density of 1.0 ×10^5^ cells/mL and allowed to recover overnight. 24 h after seeding the cells were stained using imaging medium (FluoroBrite DMEM, catalog A1896701, Thermo Fisher Scientific) supplemented with 5 µg/mL Hoechst 33342 dye (Catalog 62249, Thermo Fisher) and 1x Cell Brite Red (Catalog 30108-T, Biotinum) for nucleus and membrane staining. After staining, cells were imaged as non-stressed in isosmotic imaging medium (FluoroBrite DMEM, catalog A1896701, Thermo Fisher Scientific) or stressed in hyperosmotic imaging medium (FluoroBrite DMEM, catalog A1896701, Thermo Fisher Scientific) supplemented with 0.5 M sorbitol.

During the observation, live cells were imaged in a stage incubator with 5% CO2 at 37°C.

Images were acquired using a Zeiss 980 Laser Scanning Confocal microscope equipped with a Plan-Apochromat 63x oil objective, 40x multi-immersion LD LCI Plan-Apochromat objective, and a 20x air Plan-Apochromat objective, LSM 980 (Zeiss Instruments). Data acquisition used ZEN 3.1 Blue software (Zeiss Instruments). Hoechst 33342 dye was excited by the 405 nm laser light with the spectral detector set to 409-481 nm. mVenus protein was excited by the 488 nm laser light with the spectral detector set to 495-550 nm.. Cell Brite Red was excited by the 639 nm laser with the spectral detector set to 640-720 nm. Images were processed using ZEN 3.1 Blue software airyscan tool. Data analysis of the Z-stacks to generate maximum intensity projections was performed in FIJI (V. 2.3.0/1.52q).

### Quantification of condensation

Images used for condensate quantification were obtained and processed as detailed above. Images were then loaded into FIJI (V. 2.3.0/1.52q). Color channels were split and the channel corresponding to 488 nm (green) was selected. To mask condensates, thresholding was performed by selecting a range of intensities from 230 to 250. The Analyze Particles tool was used on masked condensates and integrated densities recorded for the entire image.

### Cell viability assays

Cells expressing mVenus, mVenus:CAHS D, mVenus:FL_Proline and mVenus:2X Linker were seeded in two sets of triplicates at a density of 1.0 x10^5^ cells/mL in 96-well plates (Catalog 3596, Costar, Fisher Scientific).

24 h after seeding, one set of triplicates of each variant was exposed for 24 h to hyperosmotic DMEM media containing 0.5 M sorbitol (stressed cells), while the other set of triplicates was fluid changed with isosmotic DMEM media (non-stressed control cells). Cells were incubated at 37 °C under 5% CO2 atmosphere.

After 24 h of hyperosmotic stress each well was stained with 5 µg/mL Hoechst 33342 dye (Catalog 62249, Thermo Fisher), 1 µg/mL Propidium Iodide (PI) (Catalog P3566, Thermo Fisher), and 4 µM Calcein AM (Catalog C1430, Thermo Fisher) for 30 min and their fluorescence measured with a plate reader (Tecan Spark600 Cyto). Cell viability was measured using two methods for the same well, one using Hoechst 33342/PI dye set [20,107] and the other using Calcein AM/PI dye set [107]. For the Hoescht 33342/PI set measurement, Hoechst 33342-positive cells and PI-negative cells were counted as alive cells and Hoechst 33342-positive and PI-positive cells were counted as dead cells. For the Calcein AM/PI set measurement, Calcein AM-positive cells and PI negative cells were counted as alive cells and Calcein AM negative cells and PI-positive cells were counted as dead cells. The survival rates were calculated by normalizing the cells in stressed wells to the cells in non-stressed wells.

Measuring the viability of the cells using two different methods gave us a more accurate reading of the cell viability within the wells that help us avoid possible bias introduced using only one method. In all the experiments the normalized survival measured with both methods gave us the same results (Fig. S6C-E).

To measure the survival of the cells after 24 h hyperosmotic stress followed by 24 h recovery the same approach was used, but after the 24 h hyperosmotic stress both sets of triplicates were fluid changed to isosmotic DMEM and the cells were left to recover from hyperosmosis at 37 °C under 5% CO2 atmosphere for 24 h. After the recovery time the viability of the cells was measured using both methods as described above.

### ER stress and Unfolded Protein Response (UPR) assay

To assess ER stress/UPR cells were assayed using a Green/Red Ratiometric Cell Stress Assay Kit (Catalog U0901G, Montana Molecular) [60].

Following the commercial kit protocol for adherent cells, cells were seeded at 1.0 x10^5^ cells/mL in 100 µl of DMEM media in a 96 well plate. Each cell line (naive HEK293, mVenus:CAHS D, mVenus:FL_Proline, mVenus:2X Linker and mVenus) was seeded in triplicate.

For each transduction 10 µl of the Bacman sensor was mixed with 0.3 µl of 500 mM Sodium Butyrate and 39.7 µl of media and added on top of the seeded cells. Cells were incubated at 37 °C under 5% CO_2_ atmosphere.

After 4 h incubation time, media was replaced with new media supplemented with 2 mM Sodium Butyrate and cells were further incubated at 37 °C under 5% CO_2_ atmosphere for 20 h.

Media was again replaced with fluorobrite imaging media supplemented with 2 mM Sodium Butyrate and red and green fluorescence was measured in a plate reader (Tecan Spark600 Cyto), this measurement was T= 0 h. Cells expressing the sensor will have nuclear red fluorescence and cells showing ER stress/ UPR will have red and green nuclear fluorescence. Measurements were taken after 6 h (T= 6 h) and after 24 h (T= 24 h).

Background red and green fluorescence was subtracted from control cells that were not transduced. The normalized fraction of ER stress/UPR was measured dividing the green cell fluorescence by the red cell fluorescence.

### Volumetric measurements

Cells were seeded and stained as mentioned in the fluorescence imaging section. Full Z-stacks of entire cells in isosmotic imaging media or hyperosmotic imaging media were acquired using an LSM 980 (Zeiss Instruments) microscope with ZEN 3.1 Blue software (Zeiss Instruments) in airyscan to perform super resolution imaging. Individual cell volumes of stressed and non-stressed cells of all the variants were measured using Imaris Cell Imaging Software (Oxford Instruments). Images from three independent experiments were analyzed and volumes of at least 20 cells per construct and condition were measured for a total of 213 cells. To obtain the relative volume change, the cell volumes of stressed cells were normalized to the non-stressed cell volumes, and the difference was plotted as % of cell reduction.

### Time-lapse imaging of recovery from hyperosmotic stress

Cell lines expressing mV:CAHS D and mV:2X Linker variants were seeded and stained with 5 µg/mL Hoechst 33342 dye (Catalog 62249, Thermo Fisher) for nuclear staining and 1x Cell Brite Red (Catalog 30108-T, Biotinum) for membrane staining as mentioned in cell line generation and imaging section. The cells were initially observed in isosmotic imaging medium for ∼5 minutes. Then, 0.5 M sorbitol was added to the imaging medium and the cells were incubated in hyperosmosis for about 10-20 min until clear condensation of the cells was observed. Once condensation was observed, the medium was replaced again with isosmotic imaging medium, and the cells were imaged as a time lapse measuring every 5 seconds. Images were processed using ZEN 3.1 Blue software airyscan tool and data analysis was performed in FIJI (V. 2.3.0/1.52q).

### Measurement of the metabolic rate of the cells

The metabolic activity of the cells was evaluated using also two methods to validate the metabolic rate. In the first method, we measured the resorufin fluorescence using presto blue HS assay (Catalog P50200, Thermo Fisher) [108,109]. This assay measures the ability of metabolically-active cells to enzymatically reduce resazurin to resorufin. For the second method, we used Cyquant XTT (Catalog X12223, Thermo Fisher) assay. XTT is a colorimetric assay based on the reduction of a yellow tetrazolium salt (sodium 3’-[1- (phenylamino carbonyl)- 3,4- tetrazolium]-bis (4-methoxy6-nitro) benzene sulfonic acid hydrate) to an orange formazan dye by metabolically active cells. The formazan dye formed is soluble in aqueous solutions and can be directly quantified using a plate reader [109,110]. Cells expressing mVenus, mVenus:CAHS D, mVenus:FL_Proline and mVenus:2X Linker were seeded in two sets of triplicates at a density of 1.0 x10^5^ cells/mL in 96-well plates.

24 h after seeding, one set of triplicates of each variant was exposed for 24 h to hyperosmotic DMEM media containing 0.5M sorbitol (stressed cells) while the other set of triplicates was fluid changed with isosmotic DMEM media (non-stressed control cells). Cells were incubated at 37°C under 5% CO2 atmosphere.

After 24 h hours of hyperosmotic stress, either 10 µl/well of Presto blue HS or 70 µl/well of Cyquant XTT working solution was added and the cells were incubated for a fixed time of 4h in all cases. Presto Blue’s resorufin fluorescence was measured with a plate reader (Tecan Spark600 Cyto) following manufacturer instructions using an excitation/emission wavelength of 560/590 nm [108]. Cyquant XTT absorbance was measured following manufacturer instructions at 450 nm and 660 nm. Blank wells with only media were added for each assay to subtract background signal. Survival by metabolic activity was quantified, normalizing the fluorescence or absorbance of stressed cells with non-stressed cells.

The same approach was used to measure the survival by metabolic activity of the cells after 24 h hyperosmotic stress followed by 24 h recovery. After the 24 h hyperosmotic stress both sets of triplicates were fluid changed to isosmotic DMEM and cells were left to recover at 37°C under 5% CO2 atmosphere for 24 h. After the recovery time either presto Blue HS or Cyquant XTT were added and the survival by metabolic activity was measured.

To calculate the metabolic rate per live cell (Fig 6E, Fig. S6H for Presto Blue, Fig. S6I & J for Cyquant XTT), the survival by metabolic activity measured with Presto Blue HS and Cyquant XTT was normalized to the survival measured by the exclusion dyes Hoechst 33342, Calcein AM and PI (Fig. S6E).

To compare the metabolic rate to cells that don’t express any of our variants, ‘control cells’, we normalized the metabolic rates of the cells expressing CAHS D and the variants to the metabolic rates of cells expressing mVenus control (Fig. 6I for Presto Blue & Fig. S6J for Cyquant XTT).

### Quantification and statistical analysis

All-atom simulations were run using the ABSINTH implicit solvent model and CAMPARI Monte Carlo simulation engine (https://campari.sourceforge.net/). Simulations were analyzed using SOURSOP (https://github.com/holehouse-lab/soursop) and MDTraj. IDR kappa values were calculated using sparrow.

SAXS data were reduced at the beamline using the BioXTAS RAW 1.4.0 software (Hopkins et al., 2017). Buffer subtraction, subsequent Guinier fits, and Kratky transformations were done using custom MATLAB (Mathworks, Portola Valley, CA) scripts.

FTIR data was analyzed using the i-signal program (version 2.72) included in the SPECTRUM suite (http://terpconnect.umd.edu/~toh/spectrum/iSignal.html) written in MATLAB language.

The statistical analysis of cell assays was performed using Microsoft Excel (v.2201, build 14827.20198) and R studio (R v4.1.0, R studio v 1.4.1717, (packages ggplot2, gridExtra, readxl, dplyr, ggsignif). The statistical details of each experiment can be found embedded in the results section and in the figure legends, including the type of statistical test used, and significance thresholds.

Correlation plots were analyzed by generating a Pearson correlation coefficient in R Studio (R v 4.2.3, R studio v 2023.03.0+386). The subsequent p-value was used to assess the significance of each correlation.

## Acknowledgements

This work was supported by DARPA award W911NF-19-2-0019, an Institutional Development Award (IDeA) from NIH grant (P20GM103432), and NASA award 80NSSC22K1629 to T.C.B.; NSF Integrative Biology award 2128069 to S.S., T.C.B. and A.S.H. supported this study.; NIH award R35GM137926 to S.S.; Financial support from MIUR of Italy (RFO2019) is gratefully acknowledged by M.M., F.F. and G.V.; Fellowships to S.B. and J.F.R. funded by Wyoming NASA EPSCoR, NASA Grant #80NSSC19M0061 supported this work.; A.V.R. is funded by the Leverhulme Trust through a grant to D.N.W. and Dr. JJ McManus (RGP-2021-049).; A.V.R. and D.N.W. are members of University of Bristol-funded Max Planck-Bristol Centre for Minimal Biology.; E.T.U is a W.M. Keck Fellow.; This research used resources of the Advanced Photon Source, a U.S. Department of Energy (DOE) Office of Science User Facility operated for the DOE Office of Science by Argonne National Laboratory under Contract No. DE-AC02-06CH11357 and resources supported by grant 9 P41 GM103622 from the National Institute of General Medical Sciences of the National Institutes of Health. Use of the Pilatus 3 1M detector was provided by grant 1S10OD018090-01 from NIGMS.; We thank members of the Water and Life Interface Institute (WALII), supported by NSF DBI grant # 2213983, for helpful discussions.; Drs. Gary J. Pielak and Samantha Piszkiewicz (University of North Carolina, Chapel Hill, NC, USA) are acknowledged and thanked for their early discussions and efforts, which were made possible with support from NIH award R01GM127291 to G.J.P.; Lorena Rebecchi (University of Modena and Reggio Emilia, UNIMORE, Italy) is thanked for stimulating discussions and valuable advice.; G.G is grateful for the support by the Netherlands Organisation for Scientific Research (NWO), Project Number VI.Veni.212.240.

## Author Contributions

**S.S.M.** Data curation, Formal analysis, Investigation, Methodology, Supervision, Visualization, Writing - original draft, Writing - review & editing

**K.N.** Data curation, Formal analysis, Investigation, Visualization, Writing - original draft, Writing - review & editing

**S.B.** Investigation, Visualization, Writing - original draft, Writing - review & editing

**V.N.** Data curation, Formal analysis, Investigation, Visualization, Writing - original draft, Writing - review & editing

**A.V.R.** Data curation, Formal analysis, Investigation, Visualization, Writing - review & editing

**J.R.** Data curation, Formal analysis, Investigation, Visualization, Writing - original draft, Writing - review & editing

**S.K.** Data curation, Formal analysis, Investigation, Visualization, Writing - original draft, Writing - review & editing

**A.A.** Investigation, Writing - review & editing

**C.C.** Investigation, Writing - review & editing

**E.T.U.** Data curation, Formal analysis, Investigation, Visualization, Writing - review & editing

**G.M.G.** Data curation, Formal analysis, Investigation, Visualization, Writing - review & editing

**F.Y.** Data curation, Formal analysis, Investigation, Writing - review & editing

**E.G.** Data curation, Formal analysis,Investigation, Writing - review & editing

**M.M.** Data curation, Formal analysis, Investigation, Writing - review & editing

**F.F.** Data curation, Formal analysis, Investigation, Writing - review & editing

**G.V.** Data curation, Formal analysis, Investigation, Writing - review & editing

**E.W.M.** Data curation, Formal analysis, Investigation, Methodology, Writing - review & editing

**F.C.** Investigation, Writing - review & editing

**G.G.** Data curation, Formal analysis, Investigation, Visualization, Writing - review & editing

**S.W.** Methodology, Supervision, Writing - review & editing

**S.S.** Methodology, Supervision, Writing - review & editing

**D.N.W.** Methodology, Supervision, Writing - review & editing

**A.S.H**. Methodology, Supervision, Writing - review & editing

**T.C.B.** Conceptualization, Methodology, Writing - original draft, Writing - review & editing

## Supplementary Text and Figures

To determine what ensemble structure within the termini drives gel formation, we performed femtosecond two-dimensional infrared (2D-IR) spectroscopy on CAHS D solutions at concentrations below and above the critical concentration for gelation. The amide I vibration of the backbone amide groups is particularly sensitive to the protein conformation [103]. In particular, in β-sheet and α-helix structures, the spatial arrangement of the amide groups causes excitonic couplings between the amide I vibrations, thus giving rise to delocalized normal modes, which in the case of β-sheet absorbs at 1620–1630 and 1680–1700 cm^−1^ [103]. 2D-IR Spectroscopy detects the couplings between these normal modes, which appear as off-diagonal features (cross peaks) in the two-dimensional spectrum [111]. These 2D-IR cross peaks are reliable markers for the presence of β-sheet structures [47,48], in particular in the case where β- sheet features may be difficult to discern in the conventional FTIR spectrum, for instance due to the presence of other secondary structures in the protein [49]

Figure 4C shows the 2D-IR spectrum of CAHS D above the gelation threshold (2.5 wt%). We observe two intense diagonal peaks at pump frequencies of 1630 and 1650 cm^−1^, and a weaker one at 1690 cm^−1^. When exciting at the pump frequency of 1630 cm^−1^, we observe a cross-peak signature at a probe frequency of 1690 cm^−1^, which is a marker of β-sheet structure [48,49]. Figure 4D shows a horizontal slice through the 2D-IR spectrum (obtained by averaging over the pump-frequency range from 1625 to 1635 cm^−1^) in which the cross peak is better visible. The cross peak is absent in the 2D-IR signal when measuring at a CAHS D concentration of 0.5 wt% (Fig. 4D), indicating that CAHS D only adopts a β-sheet conformation when the concentration is above the critical concentration for gelation.

We then assessed how the conformation of CAHS D changes in going from the gelled to the desiccated state. To this end, we performed FTIR experiments on dried CAHS D gels by fitting the infrared spectra using three Gaussian-shaped bands in agreement with the 2D-IR results. We observed that β-sheet content increases in the hydrated state as a function of concentration (Fig. 4D). This increase continues in the drying hydrogel (100- 95% relative humidity) but then begins to decrease at lower hydration levels (75-11% relative humidity) (Fig. 4E). This implies that there is an optimal hydration level for stabilizing β-β contacts, which may relate to the need for higher stability while the matrix is undergoing the final stages of drying or the early stages of rehydration.

**Supplementary Figure 1 (Related to Figure 1).**
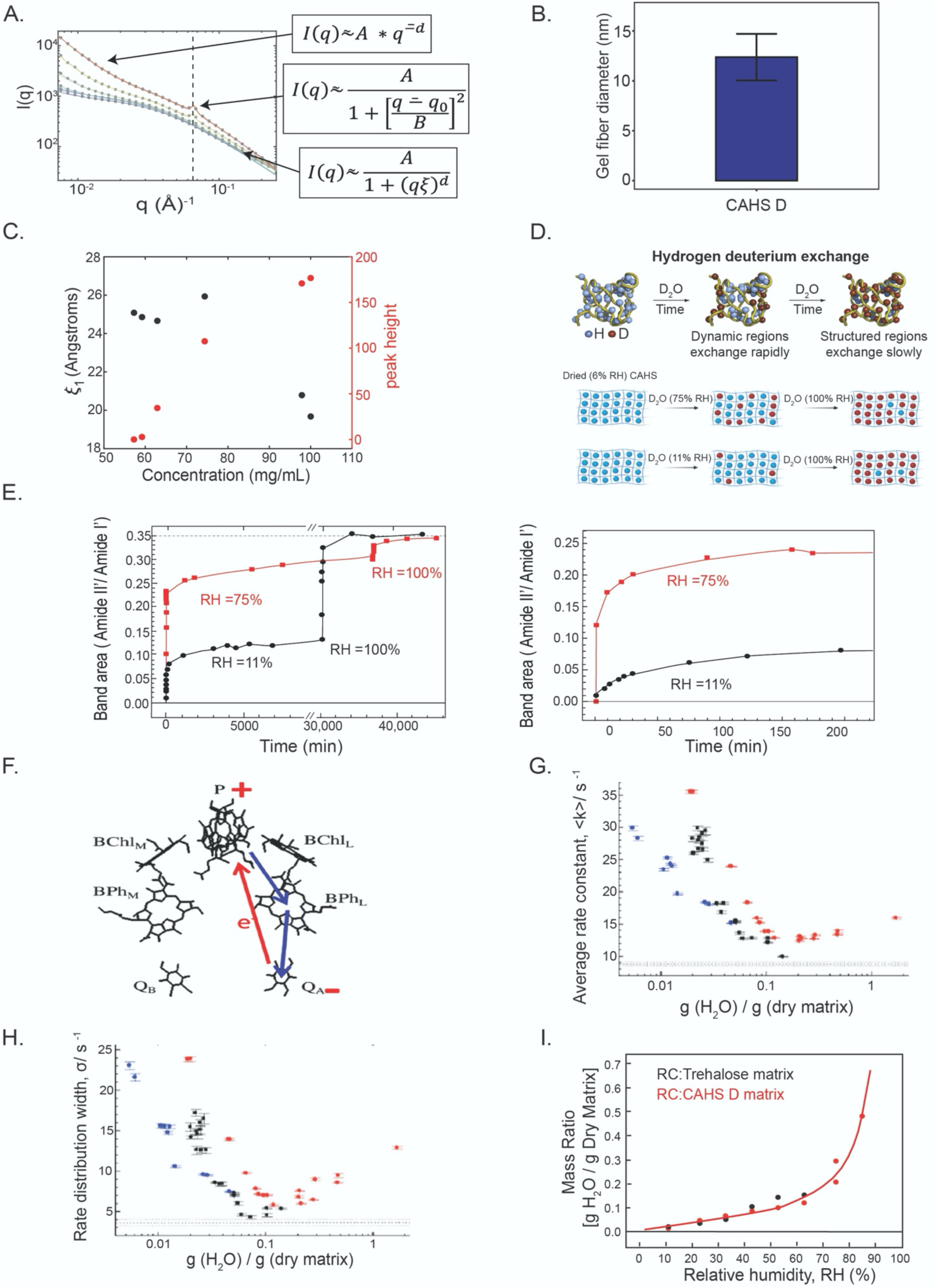
**A)** Concentration gradient SAXS on CAHS D and associated equations. **B)** Quantification of gel fiber diameter from SEM images of CAHS D (n = 20). **C)** SAXS analysis and quantification of void space sizing within a CAHS D fiber as a function of concentration of CAHS D. **D)** Schematic representation of hydrogen deuterium exchange experiment. **E)** Hydrogen deuterium exchange results for full time range (left) and for just the initial 200 minutes (right). **F)** Schematic representation of the cofactors of the bacterial photoreactive center used, involved in photoinduced electron transfer and charge recombination. **G)** Average rate constant <k> and **H)** rate distribution width σ of charge recombination kinetics of the photoreactive center (see eq. 1) as a function of hydration level for the reaction center alone (blue), reaction center in a trehalose glass (black), or the reaction center in a CAHS D gel (red). Dashed horizontal lines correspond to <k> and σ values determined in solution. **I)** Mass ratio of water to dry matrix as a function of relative humidity for a sample composed of trehalose (black) or CAHS D (red) - note both glassy matrices contain similar degrees of water content.

**Supplementary Figure 2 (Related to Figure 2).**
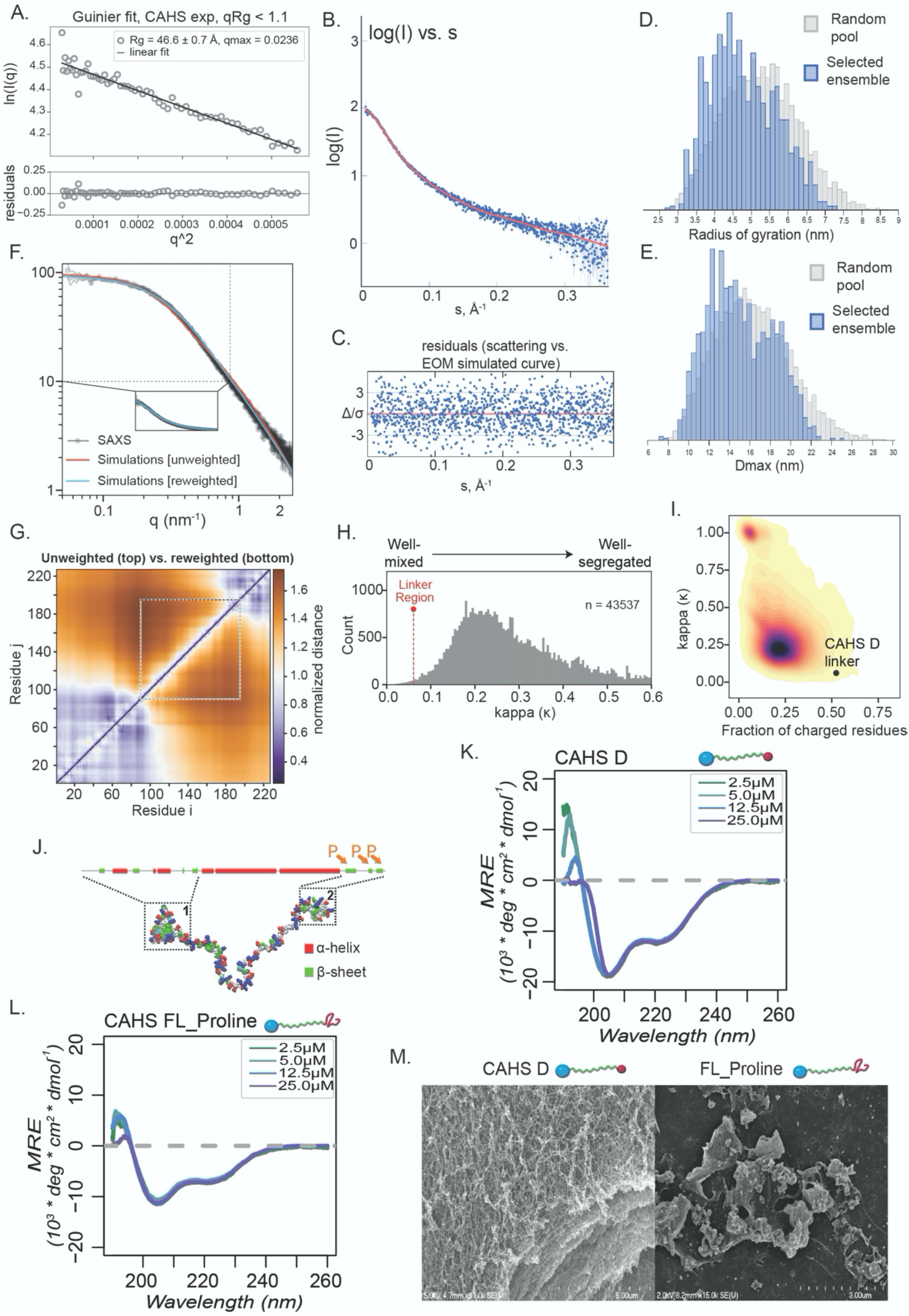
**A)** Guinier analysis of scattering data with residuals shown below. **B)** EOM-derived scattering profile compared to experimental data in log(I) vs. s space. **C)** Normalized distribution of experimental SAXS data points around the EOM-derived curve shown in panel B. **D)** Frequency of R_g_ values observed in the initial pool of structures generated by EOM and in subset of structures that were selected by EOM’s genetic algorithm to fit our experimental SAXS data. **E)** Frequency of D_Max_ values observed in random CAHS D structures and in the ensemble selected by EOM. **F)** Comparison of scattering profiles derived from re-weighted simulation ensemble vs. unweighted simulation ensemble. The difference in overall radius of gyration upon reweighting changes from 5.15 nm to 4.88 nm, suggesting this is a small change, in line with the modest changes observed with the scattering profile. **G)** Comparison of inter-residue normalized distance map for unweighted (top left) and re-weighted (bottom right) simulated ensembles confirms no major changes in intramolecular interactions occur upon reweighting. **H)** Plot showing the distribution of kappa values within all IDRs in the *H. exemplaris* proteome with the CAHS D linker annotated as being in the top 1% most well-mixed IDRs. **I)** Kernel density plot highlighting the distribution of IDRs in the *H. exemplaris* proteome in terms of fraction of charge residues and charge patterning. The CAHS D linker is highlighted as an extreme outlier in both dimensions. **J)** Schematic CAHS D with predicted secondary structure and location of Full-Length Proline (FL_Proline) prolines highlighted. **K)** Circular dichroism spectroscopy of CAHS D as a function of concentration. **L)** Circular dichroism spectroscopy of FL_Proline as a function of concentration. **M)** SEM images of FL-Proline at 50 g/L compared to CAHS D gel.

**Supplementary Figure 3 (Related to Figure 3).**
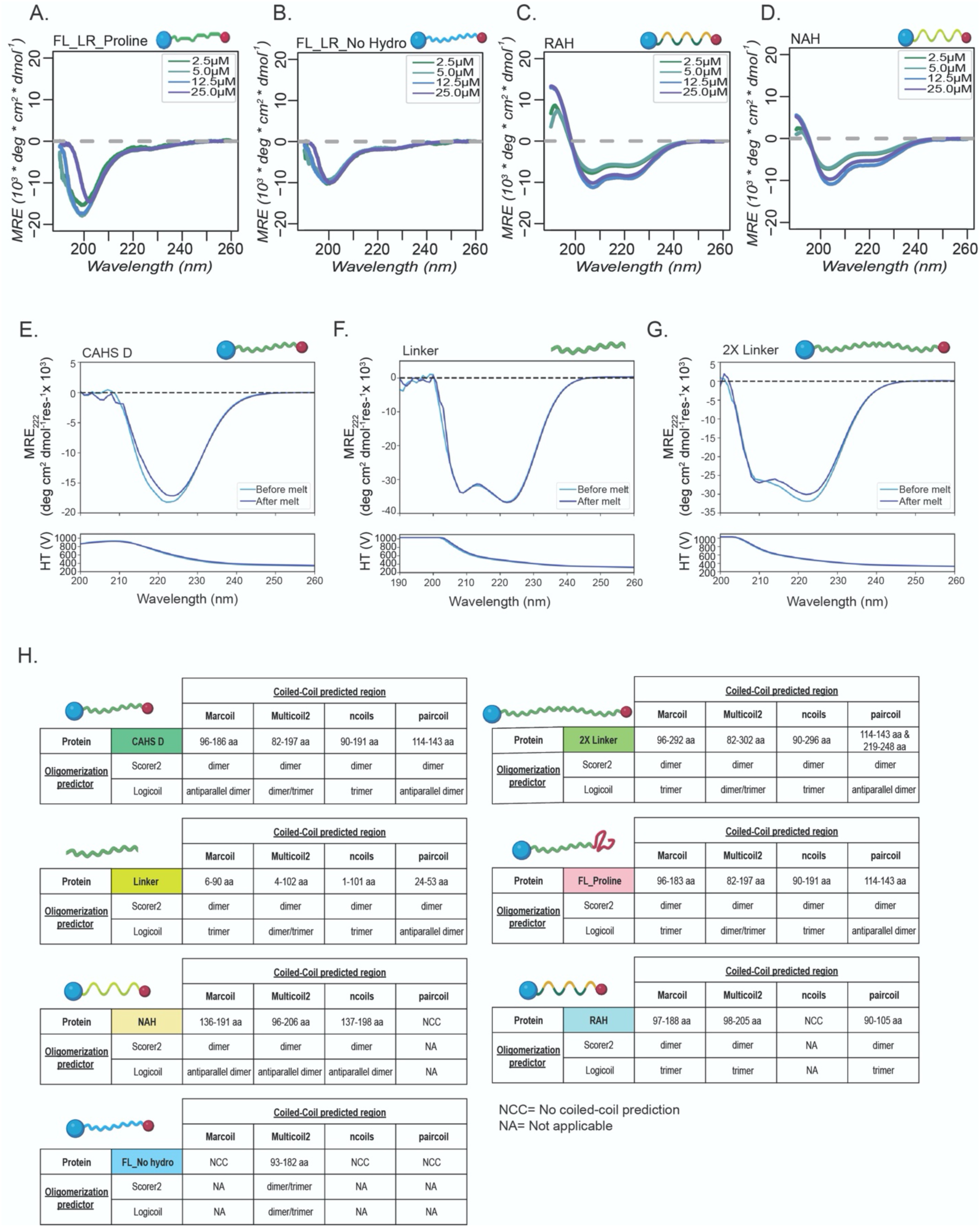
**A)** Circular dichroism spectroscopy as a function of concentration of FL_LR Proline, **B)** FL_LR No hydro, **C)** RAH and **D)** NAH variants **E)** CD spectra before (light blue line) and after (dark blue line) thermal denaturation of gelled CAHS D (0.7 mM, 17.7 mg/mL), **F)** LR variant (0.7 mM, 8.6 mg/mL) and **G)** Gelled 2X LR variant (0.25 mM, 9.3 mg/mL). **H)** Coiled-coil predictions for linker variants.

**Supplementary Figure 4 (Related to Figure 4).**
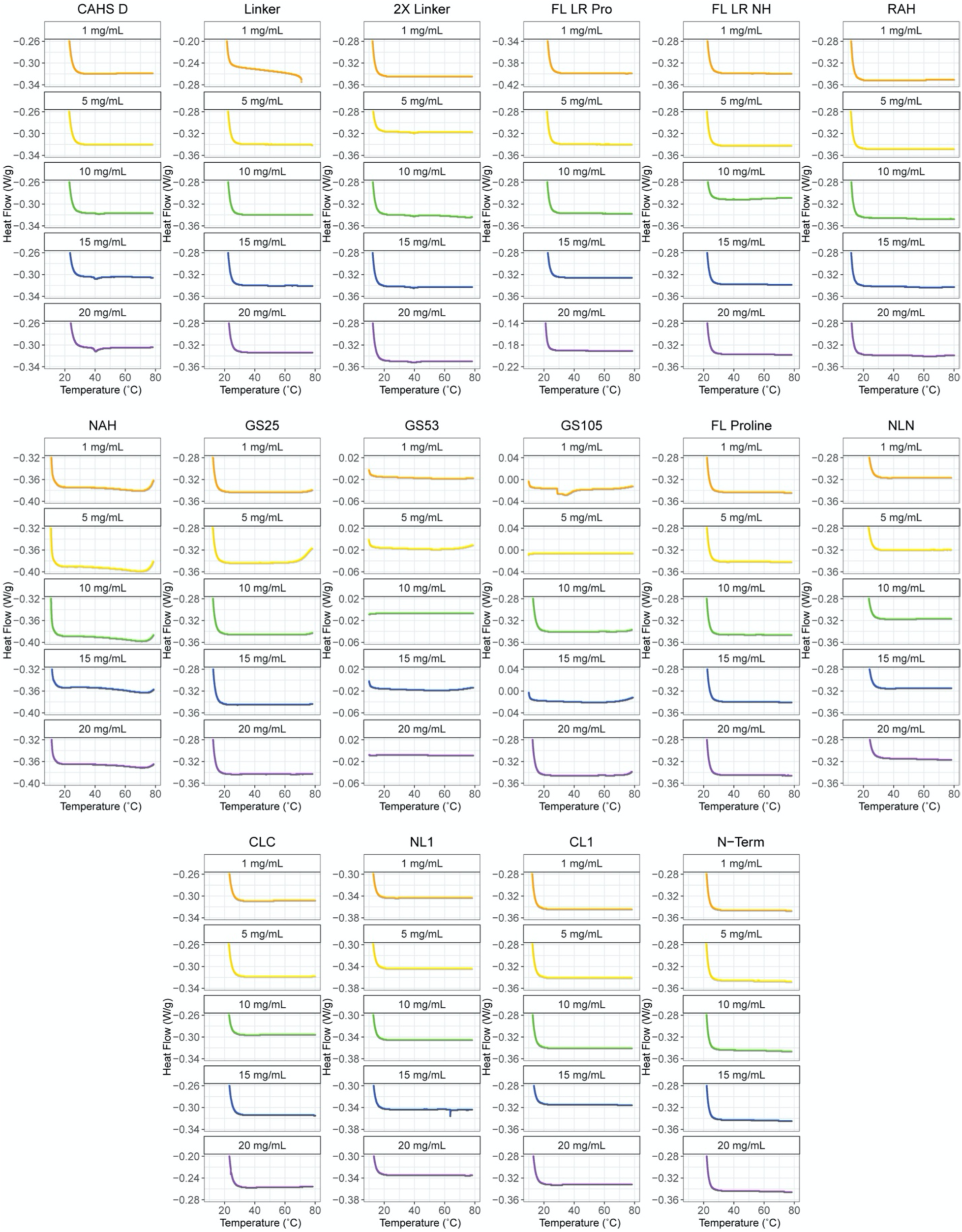
**A)** Differential scanning calorimetry thermogram displaying melt curves for CAHS D and its variants at different concentrations.

**Supplementary Figure 5 (Related to Figure 5).**
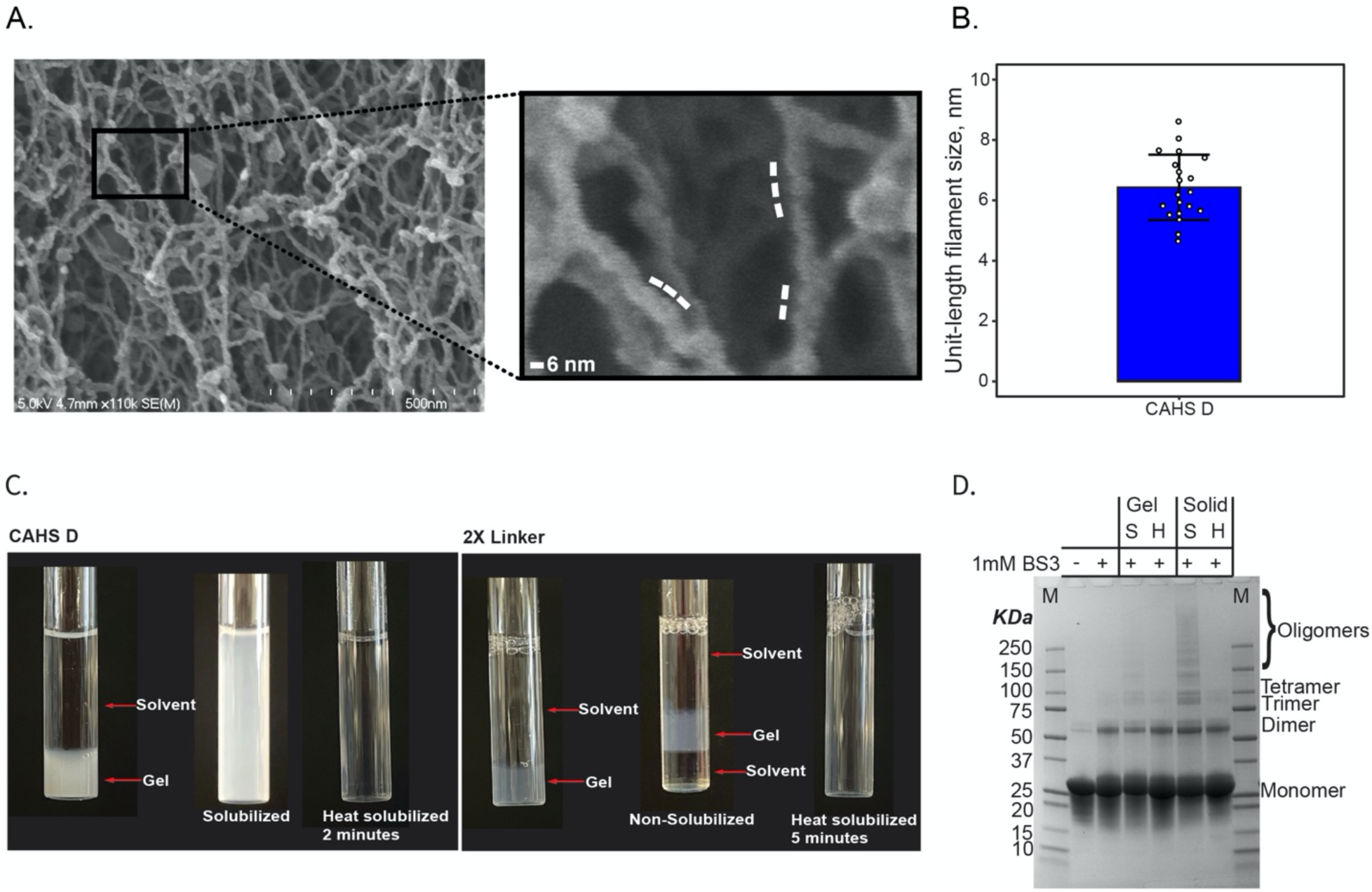
**A)** Representative SEM image of CAHS D gel (50 g/L). **B)** Quantification of “bumps” (Unit-length filaments) on CAHS D fibers. **C)** Dilution of 10 g/L (0.4 mM) CAHS D gel (left) and 2X Linker gel (right) in 20 mM Tris buffer results in instant gel resolvation for CAHS D but no gel resolvation for 2X Linker. Heat resolvation at 55°C happens within ∼2 min for CAHS D and ∼ 5 min for 2X Linker. **D)** SDS-Page showing resolvation of crosslinked CAHS D gels and solids. First lane shows MW marker. Second lane has 2 mg/mL CAHS D without crosslinker showing monomeric state. Third lane has 2 mg/mL CAHS D crosslinked with 1 mM BS3 showing formation of mostly dimers. Fourth lane shows resolvated CAHS D gel with buffer, down to 2 mg/mL concentration, crosslinked with 1 mM BS3, showing dimers and high order oligomers. Fourth lane is identical to third lane but the gel has been further heat resolubilized prior crosslinking, showing the disappearance of the high order oligomers mirroring the clear solution we see after 55°C heating in Fig. S5C. Lanes five and six are the same as third and fourth but starting the resolvation from a gel that has been dessicated in a speedvac for 16h to turn it into a crystalline solid. As in the third lane, after solubilizing with buffer there are still high order oligomers that disappear after heating (lane six).

**Supplementary Figure 6 (Related to Figure 6).**
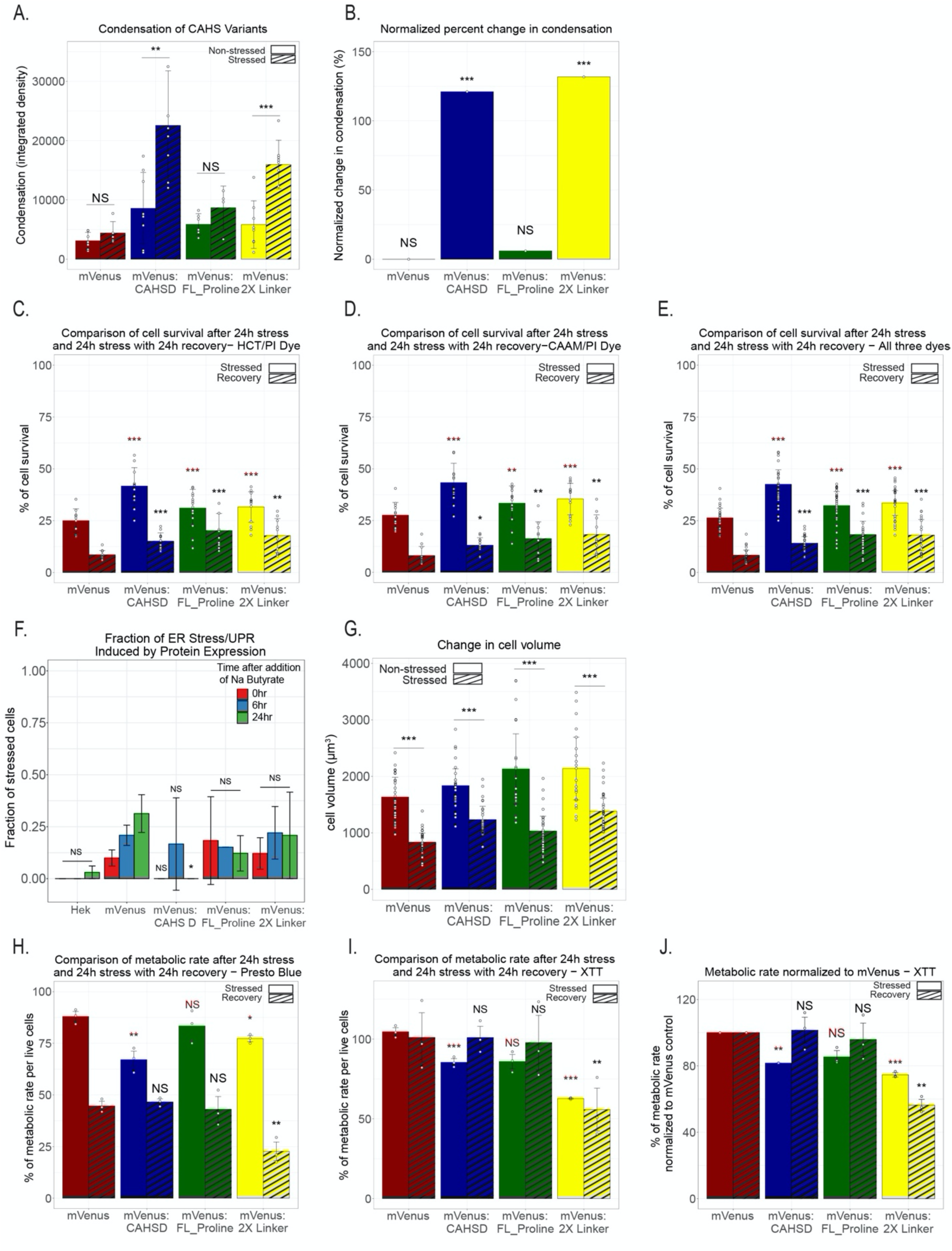
**A)** Quantification of condensation of CAHS D and its variants before (solid bars) and during osmotic shock (striped bars). **B)** Quantification of change in condensation for CAHS D and its variants during osmotic stress. **C)** Quantification of cell viability during osmotic stress (solid bars) and after recovery (striped bars) using the Hoechst/Propidium iodide assay. **D)** Quantification of cell viability during osmotic stress (solid bars) and after recovery (striped bars) using the Calcein AM/Propidium iodide assay. **E)** Quantification of combined data from Calcein AM/Propidium iodide and Hoechst/Propidium iodide viability assays during osmotic stress (solid bars) and after recovery (striped bars). **F)** ER stress and unfolded protein response from HEK cells expressing mVenus:CAHS D, mVenus:FL_Proline and mVenus:2XLinker compared to overexpressing mVenus protein and naive HEK cells. **G)** Quantification of cell volume before (solid bars) and during osmotic stress (striped bars) in cells expressing CAHS D or one of its variants. **H)** Quantification of metabolic rates of alive cells during (solid bars) and after (striped bars) osmotic stress using the Presto Blue HS assay. **I)** Quantification of metabolic rates of alive cells during (solid bars) and after (striped bars) using the XTT assay. **J)** Normalized metabolic rates of alive cells to mVenus control cells during osmotic stress (solid bars) and after recovery (striped bars) using the XTT assay. Error bars represent average deviation. Significance determined using a paired T. Test. Asterisks represent significance relative to non-stressed cells in figures A and G. In Figure B asterisks represent significance to mVenus expressing cells. In figures C-E and G-I red color statistics represent significance relative to mVenus stressed cells, and black color statistics represent significance to mVenus recovered cells. *p<0.05, **p<0.01, ***p<0.005, NS is not significant.

